# Endocrine persistence in ER+ breast cancer is accompanied by metabolic vulnerability in oxidative phosphorylation

**DOI:** 10.1101/2024.09.26.615177

**Authors:** Steven Tau, Mary D. Chamberlin, Huijuan Yang, Jonathan D. Marotti, Alyssa M. Roberts, Melissa M. Carmichael, Lauren Cressey, Christo Dragnev, Eugene Demidenko, Riley A. Hampsch, Shannon M. Soucy, Fred Kolling, Kimberley S. Samkoe, James V. Alvarez, Arminja N. Kettenbach, Todd W. Miller

## Abstract

Despite adjuvant treatment with endocrine therapies, estrogen receptor-positive (ER+) breast cancers recur in a significant proportion of patients. Recurrences are attributable to clinically undetectable endocrine-tolerant persister cancer cells that retain tumor-forming potential. Therefore, strategies targeting such persister cells may prevent recurrent disease. Using CRISPR-Cas9 genome-wide knockout screening in ER+ breast cancer cells, we identified a survival mechanism involving metabolic reprogramming with reliance upon mitochondrial respiration in endocrine-tolerant persister cells. Quantitative proteomic profiling showed reduced levels of glycolytic proteins in persisters. Metabolic tracing of glucose revealed an energy-depleted state in persisters where oxidative phosphorylation was required to generate ATP. A phase II clinical trial was conducted to evaluate changes in mitochondrial markers in primary ER+/HER2-breast tumors induced by neoadjuvant endocrine therapy (NCT04568616). In an analysis of tumor specimens from 32 patients, tumors exhibiting residual cell proliferation after aromatase inhibitor-induced estrogen deprivation with letrozole showed increased mitochondrial content. Genetic profiling and barcode lineage tracing showed that endocrine-tolerant persistence occurred stochastically without genetic predisposition. Mice bearing cell line- and patient-derived xenografts were used to measure the anti-tumor effects of mitochondrial complex I inhibition in the context of endocrine therapy. Pharmacological inhibition of complex I suppressed the tumor-forming potential of persisters and synergized with the anti-estrogen fulvestrant to induce regression of patient-derived xenografts. These findings indicate that mitochondrial metabolism is essential in endocrine-tolerant persister ER+ breast cancer cells and warrant the development of treatment strategies to leverage this vulnerability in the context of endocrine-sensitive disease.

**Statement of Significance:** Endocrine-tolerant persister cancer cells that survive endocrine therapy can cause recurrent disease. Persister cells exhibit increased energetic dependence upon mitochondria for survival and tumor re-growth potential.

## Introduction

Breast cancer mortality has decreased over the past decades, primarily due to advances in screening and treatment^1^. The majority of breast tumors express estrogen receptor α (ER), which typically reflects a degree of dependence upon estrogens for growth. Early-stage ER+/HER2-breast cancer is commonly treated with surgery followed by adjuvant endocrine therapies that antagonize ER (e.g., tamoxifen) or aromatase inhibitors that suppress estrogen biosynthesis to inhibit ER transcriptional activity. However, ∼30% of patients experience recurrence within 20 years of follow-up^2^. Since the recurrence rate is steady throughout this 20-year period, a significant proportion of events occur after the standard 5-10 years of adjuvant endocrine therapy, suggesting long periods of latency during which clinically occult cancer cells tolerate endocrine therapy. Several mechanisms of endocrine resistance have been described in recurrent ER+ breast cancer, including *ESR1* mutations that constitutively activate ER^3^. However, *ESR1* mutations are rarely found in primary tumors, indicating that these aberrations arise due to selective pressure from endocrine therapy. While mechanisms of endocrine resistance that cause recurrence are well-described, little is known about mechanisms of persistence that allow ER+ breast cancer cells to lie clinically dormant.

Drug-tolerant persister cancer cells (DTPs) are the subpopulation that remains after treatment with systemic therapy (i.e., residual disease) and have been documented in multiple cancer types^4,5^. DTPs have been detected in the accessible compartments of bone marrow and blood in ER+ breast cancer patients without overt disease after years of adjuvant endocrine therapy; the presence of DTPs portends poor prognosis^6–10^. We undertook a comprehensive approach to define the endocrine-tolerant persister cell state in ER+ breast cancer and identify vulnerabilities of persister cells to inform the development of therapeutic strategies that maintain DTPs in a dormant state or eradicate them to prevent tumor recurrence.

## Materials and Methods

### Cell culture

MCF7, T47D, HCC1428, and MDAMB415 cells were obtained from American Type Culture Collection (ATCC). Cells were maintained in DMEM supplemented with 10% FBS (HyClone Laboratories). Hormone deprivation (HD) involved culture in phenol red-free DMEM containing 10% dextran/charcoal-stripped FBS (DCC-FBS; Hyclone Laboratories) and 2 mM Glutamax (ThermoFisher Scientific). Cell lines were confirmed to be mycoplasma-free (Universal Mycoplasma Detection Kit; ATCC) and authenticated by STR genotyping (University of Vermont Cancer Center DNA Analysis Facility). For experiments involving galactose replacement of glucose, phenol red-free DMEM powder was dissolved in water and supplemented with 2 mM Glutamax, 1 mM sodium pyruvate, 44 mM sodium bicarbonate, 10% DCC-FBS, and either 10 mM glucose or galactose.

### Sulforhodamine B (SRB) assay

This viability assay was performed with modifications from ref.^11^. Cells seeded in 96-well plates were treated as specified in figure legends, and drugs/media were refreshed every 3-4 d. At the terminal time point, media were removed, cells were rinsed with PBS, and cells were fixed with 10% trichloroacetic acid at 4°C for 30 min. Wells were then washed with water and allowed to air dry. Wells were stained with 0.04% SRB (Sigma) in 1% acetic acid for 30 min. SRB stain was removed, and wells were washed 5 times with 1% acetic acid and allowed to air dry. SRB was solubilized with 10 mM Tris-HCl pH 7.5, and absorbance at 490 nm was read on a plate reader.

### Serial cell imaging and analysis

Cells were seeded in 12-well plates (10,000-50,000 cells/well). Plates were imaged every 3-4 d on a Cytation 5 (BioTek) with 4x magnification and phase contrast. Images were analyzed and cell number was quantified with Gen5 software (Biotek).

### Lentiviral transfer plasmids

The human Brunello CRISPR knockout pooled library^12^ was a gift from David Root and John Doench (Addgene #73178). Plasmid lentiCas9-Blast was a gift from Feng Zhang (Addgene #52962). To construct *NDUFC1* and *NDUFB11* knockout cell lines, CRISPR/Cas9 sgRNA sequences were extracted from the Brunello Library, and oligonucleotides were synthesized by IDT and cloned into plasmid lentiGuide-Puro (gift from Feng Zhang; Addgene #52963) as described^13^. sgRNAs were stably integrated into cell lines by lentiviral transduction (described below). To generate cells stably expressed miRFP720, miRFP720 cDNA was PCR-amplified from plasmid pmiRFP720-N1 (gift from Vladislav Verkhusha; Addgene #136560) and inserted into plasmid pLVX-M-puro (gift from Boyi Gan; Addgene #125839) by isothermal assembly. Lentiviral transfer plasmid PMXS-NDI1 was a gift from David Sabatini (Addgene #72876).

### Lentivirus generation and viral transduction

Lentiviral particles were generated using standard protocols. LentiX cells (Takara Bio) were transfected with lentiviral transfer plasmid and packaging plasmids psPAX2 and pMD2.G (gifts from Didier Trono; Addgene #12260 and #12259) using Lipofectamine 2000 (ThermoFisher Scientific). At 6 h post-transfection, medium was changed. Virus-containing medium was harvested at 24 and 48 h post-transfection, and filtered (0.45 uM) before immediate use or snap-freezing and storage at −80°C. Target cells were transduced with virus in medium containing 8 ug/mL polybrene for 6 h. Cells were allowed to recover for 48 h before selection.

### Real-time Cellular Metabolic Analysis

Cells were seeded in 96-well Seahorse XF96 plates (40,000-60,000 cells/well; Agilent) and allowed to adhere overnight. Oxygen consumption rate (OCR) and extracellular acidification rate (ECAR) were serially measured using the Seahorse XF96 Analyzer (Agilent). ATP production rates from mitochondrial OXPHOS and glycolysis were derived from OCR and ECAR measurements using the Seahorse ATP Rate Assay Kit. Mitochondrial respiration was inferred using OCR measurements using the Seahorse XF Cell Mito Stress Test Kit.

### Luciferase reporter assay

Plasmids encoding firefly luciferase driven by an estrogen-response promoter element (gift from Dorraya El-Ashry) and CMV-*Renilla* luciferase (Promega) were co-transfected into cells using Lipofectamine 2000. The following day, cells were treated ± 1 nM E2 for 24 h. Luciferase activities were measured using the Dual-Luciferase Reporter Assay System (Promega) per manufacturer instructions on a GloMax-Multi Detection System (Promega). Firefly luciferase signal was normalized to *Renilla* signal.

### Lactate assay

Cells were seeded in 96-well plates (15,000 cells/well) and treated as indicated. Media were collected after 3 d of drug treatment, diluted 1:40 in PBS, snap-frozen, and stored at −20°C. Relative lactate levels in media were measured with Lactate-Glo assay (Promega). Relative cell numbers were determined by SRB assay. Relative lactate level was normalized to relative cell number.

### ATP assay

Cells were seeded in 96-well plates (5,000 cells/well) and treated for 3 d. Media were then removed and replaced with PBS. Relative ATP levels were assessed using CellTiter-Glo 2.0 viability reagent (Promega): cells were incubated with reagent for 10 min, and lysate was then transferred to white-walled 96-well plates for luminescence measurement. Relative cell numbers were determined by SRB assay. Relative ATP level was normalized to relative cell number.

### Antibodies

Primary antibodies used included Alexa Fluor594-conjugated anti-Tom20 (Santa Cruz Biotechnology 17764), Ki67 (Biocare Medical, CRM325B), cleaved caspase 3 (Cell Signaling Technology 9505), geminin (Cell Signaling Technology 52508), ERα (Dako SP1), MT-CO2 (Sigma Aldrich HPA051505), OXPHOS antibody cocktail (Abcam ab110411), c-Myc (Cell Signaling Technology 5605), cyclin D1 (Cell Signaling Technology 2978), phospho-histone H3 Ser10 (Cell Signaling Technology 9701), HIF1α (Genetex GTX127309), CPT1α (Abcam ab128568), p21 (Cell Signaling Technology 2947), p27 (Cell Signaling Technology 3686), NDUFB11 (Proteintech 16720-1-AP), NDUFC1 (Proteintech 23842-1-AP), β-actin (Cell Signaling Technology 3700), and vinculin (Cell Signaling Technology 13901). Secondary antibodies included HRP-conjugated anti-mouse (Cytiva NA931V) and HRP-conjugated anti-rabbit (Cytiva NA9340V).

### Flow cytometry

Cells were trypsinized and resuspended in serum-containing medium with either 100 nM MitoTracker Deep Red, 100 nM TMRE, 2 uM JC-1, 100 uL/mL 2-NBDG, or 2.5 uM Nonyl Acridine Orange (NAO). For antibody staining, cells were fixed in 4% formaldehyde in PBS for 15 min, washed with PBS, and permeabilized with an ice-cold mixture of 90% methanol and 10% PBS for 30 min. Cells were re-suspended in flow cytometry buffer (0.5% BSA, 0.1% sodium azide in PBS) containing Alexa Fluor594-conjugated anti-Tom20 (1:100) and incubated for 1 h at room temperature in the dark. Cells were washed and re-suspended in flow cytometry buffer. For cell cycle analysis, cells were fixed in 70% ethanol at −20°C overnight, washed with PBS, and stained with propidium iodide/RNase staining solution (ThermoFisher Scientific) for 30 min at room temperature in the dark. Samples were analyzed on a MACSQuant-10 (Miltenyi Biotec) or a ZE5 Cell Analyzer (Bio-Rad). Data were analyzed using FlowJo 10.8.2 software (BD Biosciences). Cell cycle analysis was performed using the Cell Cycle platform within FlowJo with the Watson Pragmatic algorithm.

### Fluorescence-activated cell sorting (FACS) for mitochondrial content

Cells were serum-starved for 2 d to drive cell cycle synchronization in G1. Cells were labeled with 1 nM MitoTracker Deep Red, and resuspended in growth medium. MitoTracker^Dim^ and MitoTracker^Bright^ cells were collected as the 15% of cells with the respective lowest and highest signal intensity using a SH800 sorter (Sony). Collected cells were then reseeded in 96-well plates in medium containing penicillin/streptomycin and used for viability assay as indicated in figure legend.

### Immunoblotting

Cells were lysed in RIPA buffer (20 mM Tris, pH 7.4, 150 mM NaCl, 1% NP-40, 10% glycerol, 1 mM EDTA, 1 mM EGTA, 5 mM NaPPi, 50 mM NaF, 10 mM Na β-glycerophosphate) with HALT protease inhibitor cocktail and 1 mM Na_3_VO_4_. Lysates were sonicated for 10 sec and centrifuged at 17,000 x *g* for 10 min at 4°C. Supernatant was collected, and protein concentration was measured by BCA assay (Pierce). Cell lysates were reduced and denatured using NuPAGE (ThermoFisher Scientific) plus 1.25% β-mercaptoethanol. Fifty ug of protein/sample was analyzed by SDS-PAGE. Protein was transferred onto nitrocellulose membrane and blocked with 5% BSA in TBS containing 0.1% Tween-20 (TBS-T) for 1 h. Membranes were incubated with primary antibody overnight at 4°C on a shaker in blocking solution. Membranes were then washed with TBST 3 times for 10 min, and incubated with HRP-conjugated secondary antibody in 5% milk in TBS-T for 1 h. After 3 washes with TBS-T for 10 min each, signal was developed using Pierce ECL Western Blotting Substrate or Supersignal West Pico PLUS substrate, and blots were imaged using a Chemidoc MP (Bio-Rad).

### Animal Studies

Animal studies were approved by the Dartmouth College IACUC (protocol 00002144). Female NOD/SCID/IL2Rγ^−/−^ (NSG) mice were obtained from the Dartmouth Cancer Center Mouse Modeling Shared Resource. Mice were ovariectomized and orthotopically injected with MCF7/miRFP720 cells bilaterally in inguinal mammary fat pads. An E2 pellet (1 mg in beeswax^14^) was s.c. implanted to induce tumor growth. Tumor dimensions were measured twice weekly using calipers [volume = (length x width^2^/2)]. When a MCF7/miRFP720 tumor reached 200 mm^3^, the E2 pellet was removed to induce estrogen deprivation for 60 d. *In vivo* fluorescence imaging on a Pearl Impulse with an excitation/emission of 685/720 nm was performed on 3 consecutive days for baseline measurements prior to drug treatment, and then performed weekly thereafter. HCI-017 and HCI-003 PDX tumor models were obtained from the University of Utah^15^. PDX tumor fragments (∼8-mm^3^) were orthotopically implanted bilaterally into intact (non-ovariectomized) mice. When HCI-017 and HCI-003 tumors reached 150 mm^3^, mice were randomized to drug treatments. IACS-010759 was dissolved in DMSO at 15 mg/mL, diluted in a 0.5% methylcellulose suspension, and administered by oral gavage in 100 uL at 2.5 mg/kg/day for 5 consecutive days each week. Fulvestrant was dissolved in ethanol at 500 mg/mL, diluted in castor oil to give a working concentration of 50 mg/mL, and administered s.c. in 100 uL at 5 mg/wk.

### Clinical trial

This clinical study was approved by the Dartmouth Health Human Research Protection Program, registered on clinicaltrials.gov (NCT04568616), and conducted in accordance with ICH Good Clinical Practice. The study was monitored by the Dartmouth Cancer Center Data, Safety Monitoring, and Accrual Committee. Written informed consent was obtained from all participants prior to performing study-related procedures. All procedures performed involving human participants were in accordance with the 1964 Helsinki declaration and its later amendments or comparable ethical standards. The trial protocol is appended. Objectives of this trial were to determine whether residual cancer cells exhibit altered expression of CPT1α or mitochondrial markers (compared to baseline) following neoadjuvant endocrine therapy. Post-menopausal patients with Stage I-III breast cancer without prior endocrine therapy were eligible if a tumor was ≥1 cm in diameter, and a diagnostic biopsy specimen was HER2-negative (IHC score of 0-1+, or FISH ratio of <2 if IHC was 2+ or not performed) and strongly ER+ (≥50% malignant cells positive). Patients were treated with neoadjuvant letrozole (2.5 mg/d orally) for up to 4 months prior to surgical tumor resection. Baseline (diagnostic) and post-letrozole (surgical) tumor tissue specimens were used for IHC for MT-CO2, CPT1α, and Ki67. Ki67 was scored as the proportion of positively stained tumor cells using the Ki67 scoring app (unweighted global score) as recommended by the International Ki67 in Breast Cancer Working Group^16^. MT-CO2 and CPT1α were scored using histoscoring^17^. Ki67 and CPT1α scores were normalized using the log_2_(Ki67+1) method as described^18^.

### Immunohistochemical (IHC) staining

Sections of FFPE tissue were stained using a BOND RX autostainer (Leica Biosystems). Heat-induced epitope retrieval was performed with Bond Epitope Retrieval 2 pH 9 solution for Ki67, and with Bond Epitope Retrieval 1 pH 6 solution for cleaved caspase 3, geminin, ERα, MT-CO2, and CPT1α; slides were incubated at 100°C for 40 min. Primary antibody was applied for 15-60 min at room temperature. Antibody binding was visualized using the Leica Bond Polymer Refine detection kit with DAB chromogen and hematoxylin counterstain. Stained tissues were imaged with an IX73 inverted microscope (Olympus), and 3 representative fields at 200x magnification were imaged for analysis with HALO software (Indica Labs).

### CRISPR screening

#### Cas9 stably expressing cell line

MCF7 cells were transduced with lentivirus encoding Cas9 via lentiCas9-blast. At 3 d post-infection, cells were selected with 5 ug/mL blasticidin for 10 d. Cas9 mRNA expression was confirmed by RT-qPCR.

#### CRISPR library transduction

MCF7/Cas9 cells were seeded in 12-well plates (3×10^6^ cell/well in 40 wells), spinfected with gRNA-encoding lentivirus (MOI=0.3) at 650 x *g* for 2 h at 37°C with 8 ug/mL polybrene, and incubated for 4 h at 37°C. Medium was removed, and cells were trypsinized and reseeded into twenty 150-mm dishes. At 2 d post-transduction, cells were selected with 1 ug/mL puromycin for 5 d, and then maintained for an additional 7 d. Cells were trypsinized and divided into 3 groups using 35×10^6^ cells/group to maintain 500x sgRNA coverage: Group 1 was the Day 0 baseline sample that was snap-frozen; Groups 2 and 3 were reseeded into ten 150-mm dishes each. Groups 2 and 3 were treated with or without 1 nM E2 for 3 wk, followed by trypsinization and snap-freezing.

#### gDNA extraction, processing, and sequencing

gDNA was purified using the QIAamp Blood DNA Maxi kit (Qiagen). A total of 100 ug DNA per sample was split into 100 PCR reactions (primers: GAGAGGGCCTATTTCCCATGATTC and ATTCTTTCCCCTGCACTGTACCC) with 1 ug of gDNA template for 30 cycles, and PCR products were pooled. PCR product was purified by QIAquick PCR Purification kit (Qiagen). Amplicon size was confirmed by gel electrophoresis. A subsequent PCR was performed with 100 ng DNA per sample for 10 cycles to attach Illumina adapters. PCR product was purified with a double-sided clean up by adding 0.8x and subsequently 2x KAPA pure beads. Libraries were sequenced using a NextSeq 500 (Illumina) on High Output for 150 cycles.

#### Data analysis

The 5’ ends of reads were trimmed to 5’-CACCG-3’ using Cutadapt. MAGeCK (v.0.5.9)^19^ was used to extract read counts for each sgRNA using the ‘count’ function. The ‘mle’ function was used to compare read counts from cells treated ± E2 after controlling for Day 0 baseline counts and library size. Differential β-score for each gene was calculated by taking the difference in β-score between the ± E2 conditions. Enrichment analysis was performed using STRING^20^. Mitochondria-associated genes were annotated with MitoCarta3.0^21^.

### Cell barcoding and analysis

#### Lentiviral barcoding

MCF7 cells were transduced with the Cellecta 50M Lentiviral Barcode Library^22^. To ensure adequate representation, the number of unique barcodes was maintained at 20,000^22^.

#### *In vivo* xenografting of barcoded cells and xenograft treatment

Cells were injected orthotopically bilaterally into the inguinal mammary fat pads of ovariectomized NSG mice (5×10^6^ cells per injection). An E2 pellet was simultaneously implanted s.c. An aliquot of the baseline cell population was snap-frozen for later processing. Tumors were E2-driven for 30 d, and then E2 pellets were removed. Tumors were harvested at the indicated time points and snap-frozen.

#### DNA extraction and barcode library preparation

E2-driven tumors were thawed in Buffer P1 with RNase A (Qiagen) and 0.5% SDS, and sonicated to shear DNA. DNA was purified using phenol:chloroform:isoamyl alcohol and precipitated overnight with sodium acetate, isopropanol, and 20 ug glycogen. The DNA pellet was washed with 70% ethanol, air-dried, dissolved in water, and solubilized by incubation at 80°C for 30 min. E2-deprived tumors and the baseline cell population were homogenized by bead-beating with a 1600 MiniG tissue homogenizer in Qiagen buffer RLT Plus. DNA was isolated with the AllPrep DNA/RNA Mini Kit (Qiagen).

Libraries were prepared for sequencing by a two-step PCR amplification. In total, 500 ng genomic DNA (reflecting ∼75,000 genomes) was used for each reaction, and a total of 5 ug DNA (reflecting ∼900,000 genomes) was used per sample. The first step of the reaction amplified the barcode region using custom primers for 20 cycles with NEB Q5 Hot Start DNA Polymerase (New England Biolabs). Primer sequences are listed in Supp. Table 1. The products of the first PCR reaction were pooled for each sample and cleaned up with Kapa Pure Beads at 2x concentration. A portion (5 uL) of the purified PCR product was used for the second PCR step with custom P5 and P7 indexed primers for an additional 20 cycles with NEB Q5 Hot Start DNA Polymerase. Amplicons were cleaned up with Kapa Pure Beads at 2x concentration. DNA was quantified by Nanodrop, adjusted to the same concentration across samples, and size was verified by Fragment Analyzer (Agilent). Libraries were sequenced on a MiniSeq High-Output 75-cycle run (Illumina).

#### Barcode counting and analysis

Reads were filtered for a Phred score >20 using fastq-filter^23^. A list of barcodes across all samples was generated and counted using the UMI-tools^24^ ‘whitelist’ command. A Hamming distance of 2 was allowed to correct for errors. Shannon diversity index was calculated as 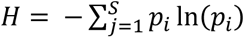, where *p_i_* is the proportion of barcode *i* ^25^.

### Quantitative Proteomics

#### Sample processing and mass spectrometry

Cells were treated as indicated, washed with PBS, scraped from dishes, centrifuged at 300 x *g* for 5 min, and pellets were snap-frozen. Cell pellets were resuspended in ice-cold lysis buffer (8 M urea, 25 mM Tris-HCl pH 8.6, 150 mM NaCl) containing phosphatase inhibitors and protease inhibitors (Roche Life Sciences) and lysed by sonication for 15 sec. Lysates were centrifuged at 15,000 x *g* for 30 min at 4°C, and supernatants were transferred to a new tube. Protein concentrations of lysates were determined using BCA assay. DTT (5 mM) and iodoacetamide (15 mM) were added to reduce and alkylate proteins, respectively. Samples were incubated overnight at 37°C with 1% trypsin. The trypsin digest was stopped by the addition of 0.25% trifluoroacetic acid (TFA). Precipitated lipids were removed by centrifugation at 3500 x *g* for 15 min, and peptides in the supernatant were desalted over an Oasis HLB 60 mg plate (Waters). An aliquot containing 100 ug of peptides was removed and labeled with Tandem-Mass-Tag (TMT) reagent (ThermoFisher Scientific). Once labeling efficiency was confirmed to be ≥95%, each reaction was quenched by the addition of hydroxylamine to reach 0.25% for 10 min, mixed, acidified with TFA to a pH of 2, and desalted over an Oasis HLB 10 mg plate (Waters). The desalted multiplex was dried by vacuum centrifugation and separated by pentafluorophenyl (PFP)-based reverse-phase HPLC fractionation as described^26^. TMT-labeled peptides were analyzed on an Orbitrap Lumos mass spectrometer (ThermoScientific) equipped with an Easy-nLC 1200 (ThermoScientific), and raw data were searched and processed as described^27^. Peptide intensities were adjusted based on total TMT reporter ion intensity in each channel.

#### Data processing

Data were analyzed using R software. Proteins with only 1 identified peptide sequence were excluded. Variance stabilizing normalization was used to normalize and transform data. UniProt IDs were converted into gene symbols; for multiple proteins mapping to 1 gene, the protein(s) with the lower variance(s) was excluded. Gene set co-regulation analysis (GESECA) was performed with the ‘fgsea’ package in R with gene set size between 15-500 genes.

### Whole-exome sequencing (WES)

Human tumor tissue specimens from 2 patients were obtained from the Dartmouth-Hitchcock Medical Center Department of Pathology tissue archive through a retrospective study approved by the Dartmouth Health Human Research Protection Program (STUDY02000731). DNA was extracted from FFPE MCF7 and human tumor tissue specimens using the QIAamp DNA FFPE Tissue kit (Qiagen). DNA was quantified using Qubit (Qiagen) and assayed by Fragment Analyzer (Agilent), followed by library preparation using the HyperPrep kit (Kapa Biosystems), multiplexing of 6-11 libraries, and exome capture using the xGen Exome kit v2 (IDT). Exomes were sequenced on an Illumina NextSeq 500. Reads were trimmed with cutadapt v.2.4^28^, and sample quality was checked with fastqc^29^. Reads mapped to the mouse genome were filtered out using xenome v1.0.0^30^. Reads were aligned to human reference genome GRCh38.p13 using BWA MEM v.0.7.17^31^. Read duplicates were flagged and excluded using Picard tools MarkDuplicates^32^ to ensure duplicated reads were not considered during variant calling. BaseRecalibrator and ApplyBQSR^32^ were used to detect and correct for patterns of systematic errors in base quality scores that could result in false-positive variant calls. To estimate copy number alterations (CNAs), CNVkit v0.9.10^33^ was performed with the default parameter. For MCF7 tumor samples, the panel of “normal samples” used were the E2-driven (baseline) samples. For human tumor samples, a flat reference was used. Segmentation was performed using the circular binary segmentation model^34^. Mutect2 from the GATK package v4.4.0.0^32^ in multi-sample mode was used to call SNVs and indels from mapped reads; a standard panel of normals was downloaded from GATK best practices Google bucket. Variants were filtered for read orientation bias using the method proposed by GATK best practices. To infer clonal composition, we employed PyClone^35^ to group sets of somatic mutations into clusters while accounting for CNAs. We integrated the variant data with CNVkit to derive total and minor copy number, and applied this to PyClone analysis. PyClone was run with default parameters and 10,000 iterations. Clusters with 1 variant or with low variant allele frequency (VAF) (VAF∼0) were considered spurious and ignored. The output from PyClone was analyzed with ClonEvol^36^ to infer clonal evolution. The assumption was made that the cluster with the highest VAF is the founder cluster. The consensus model with the highest likelihood was plotted with fishplot^37^.

### 13C-glucose tracing metabolic flux analysis

#### Treatment

MCF7 cells were treated with phenol red-free DMEM with 10% DCC-FBS ± 1 nM E2 for 21 d prior to reseeding into 10-cm dishes (5 million cells/dish). Two days later, cells were labeled with medium containing U-^13^C-glucose for 45 min. Medium was prepared by dissolving DMEM powder (without glucose, L-glutamine, phenol red, sodium pyruvate, or sodium bicarbonate) in water, and adding Glutamax (final concentration 2 mM), sodium pyruvate (final concentration 1 mM), sodium bicarbonate (final concentration 44 mM), sodium oleate (final concentration 100 uM), and U-^13^C glucose (Cayman Chemical, final concentration 25 mM). Cells were then washed with 150 mM ammonium acetate, liquid was aspirated, and liquid nitrogen was poured into the dish to snap-freeze. Dishes were placed on dry ice and stored at −80°C.

#### Sample processing and analysis

Frozen dishes were sent, processed, and analyzed at the University Michigan Metabolomics Core. Cell culture plates were removed from −80°C storage and maintained on wet ice throughout the processing steps. To each 10-cm plate, 1 mL of 80/20 methanol/water (both LC-MS grade) was added. Plates were gently agitated to release cells, and then scraped to homogenize cells and transferred to a microtube. Microtubes were vortexed and incubated at 4°C for 10 min to complete metabolite extraction.

Samples were vortexed a second time, and then centrifuged at 10,000 x *g* for 10 min at 4°C. Post-centrifugation, 200 uL of extraction solvent was transferred to an autosampler vial, then taken to dryness using a gentle stream of nitrogen at room temperature. Ten uL of extract from each sample was added to a separate autosampler vial to create a pooled sample for quality control purposes. Dried samples were reconstituted in 100 uL of 80/20 water/methanol for LC-MS analysis. Ten uL of extract was removed and pooled in a separate autosampler vial for quality control purposes.

Analysis was performed on an Agilent system consisting of an Infinity Lab II UPLC coupled with a 6545 QTOF mass spectrometer (Agilent Technologies) using a JetStream ESI source in negative mode. The following source parameters were used: Gas Temp: 250°C, Gas Flow: 13 L/min, Nebulizer: 35 psi, Sheath Gas Temp: 325°C, Sheath Gas Flow: 12 L/min, Capillary: 3500 V, Nozzle Voltage: 1500 V.

The UPLC was equipped with a 10-port valve configured to allow the column to be either eluted to the mass spectrometer or back-flushed to waste. The chromatographic separation was performed on an Agilent ZORBAX RRHD Extend 80Å C18, 2.1 × 150 mm, 1.8-um column with an Agilent ZORBAX SB-C8, 2.1 mm × 30 mm, 3.5-um guard column. The column temperature was 35°C. Mobile phase A consisted of 97:3 water/ methanol and mobile phase B was 100% methanol; both A and B contained tributylamine and glacial acetic acid at concentrations of 10 mM and 15 mM, respectively. The column was back-flushed with mobile phase C (100% acetonitrile, no additives) between injections for column cleaning.

The LC gradient was as follows: 0-2 min, 0% B; 2-12 min, linear ramp to 99% B; 12-17.5 min, 99% B. At 17.5 minutes, the 10-port valve was switched to reverse flow (back-flush) through the column, and the solvent composition changed to 99% C. From 20.5-21 min, the flow rate was ramped to 0.8 mL/min, held until 22.5 min, then reduced to 0.6 mL/min. From 22.7-23.5 min, the solvent was ramped from 99% to 0% C while flow was simultaneously ramped down from 0.6-0.4 mL/min and held until 29.4 min, at which point flow rate was returned to starting conditions at 0.25 mL/min. The 10-port valve was returned to restore forward flow through the column at 28.5 min. An isocratic pump was used to introduce reference mass solution through the reference nebulizer for dynamic mass correction. Total run time was 30 min. The injection volume was 10 uL.

#### Data analysis

Metabolites were identified by matching the retention time and mass (± 10 ppm) to authentic standards using Agilent’s Profinder Isotopologue extraction program, which performs automatic isotope subtraction based on the expected isotope distribution for the molecular formula provided.

### Single-cell RNA sequencing (scRNA-seq)

#### Sample processing

MCF7/miRFP720 tumors were harvested from mice, minced using a razor blade, and enzymatically digested at 37°C in DMEM containing 1 mg/mL collagenase type II and 2 mg/mL DNase I with stirring for 1 h. Following digestion, tissue was passed through a 70-um filter. Dissociated cells were centrifuged at 500 x *g* for 5 min at 4°C and resuspended in ACK lysis buffer for 5 min on ice to lyse red blood cells. After washing with PBS, cell pellets were resuspended in DMEM + 10% FBS. Cells were sorted for miRFP720 positivity by FACS (Sony SH800). Cell viability and number were then assessed by acridine orange/propidium iodide staining using a Cellometer K2 cell counter (Nexcelom Biosciences).

#### Cell Capture and Library Preparation

scRNA-seq libraries were prepared following the protocol provided by the HIVE CLX scRNAseq Sample Capture User Protocol (v.1) and the HIVE CLX scRNAseq Transcriptome Recovery & Library Preparation (v.1) (Honeycomb Biotechnologies). Briefly, cells were captured on the HIVE Collector by centrifuging cells into the Collector’s microwells pre-loaded with barcoded beads. After washing the HIVE and adding the CLX Cell Preservation Solution, HIVE Collectors were stored at −80°C.

HIVE Collectors were thawed and washed with CLX Storage Wash Solution, and wells were sealed with a porous membrane remove buffers. The capture of RNA onto beads was done by adding lysis buffer onto the HIVE assembly membrane followed by CLX hybridization buffer. After removal of the membrane, CLX Bead Recovery Solution was added, and HIVE assemblies were centrifuged at 3,000 x *g* for 5 min to recover beads. Captured RNA was used for first-strand synthesis to generate cDNA followed by removal of the RNA strand to allow second-strand synthesis with random primers, yielding double-stranded cDNA with cell-specific barcodes. Samples were used for whole-transcriptome amplification with SPRI bead clean-up. Custom HIVE index primers containing P5 and P7 sequences were attached with subsequent SPRI bead clean-up. Libraries were sequenced on an Illumina NextSeq 2000 with 100 cycles of paired-end sequencing.

#### Data analysis

The output bcl2 file was converted into FASTQ format with bcl2fastq v.2.20.0.422, and further processed with Beenet v.1 to produce count tables. The number of barcodes in Beenet was set to 30,000 to capture all cell-containing barcodes. Count tables were input into Seurat v.4.3.0^38^ and merged to form a single Seurat object. Features with <300 genes, <600 unique transcript molecules, and >20% mitochondrial genes present were removed. SCTransform was used for normalization and variance stabilization. Clustering was performed in Seurat with default parameters. Differentially-expressed genes were identified with the Seurat ‘FindMarkers’ command using a Wilcoxon statistical test. Genes with adjusted *p*≤0.05 were considered significant.

### Analysis of human breast tumor gene expression datasets

Gene expression profiles of paired human ER+ breast tumor specimens acquired before and during neoadjuvant aromatase inhibitor therapy were downloaded from the NCBI Gene Expression Omnibus (GEO) (GSE20181^39^ and GSE111563^40^). Series matrix files containing log_2_-transformed normalized data were collapsed to give one (most variable) probe set per gene using GenePattern. Data were then transformed into z-scores. To determine the similarity of a tumor transcriptional state to the IACS transcriptional signature, we performed a correlation analysis to generate an R-value for each tumor^41^. R-values were compared between time points by paired t-test. Subset analyses were similarly performed using the non-cell cycle-related gene signature of IACS response (gene sets included in Suppl. Table 2).

### DepMap data analysis

DepMap data from the 22Q1 release were downloaded from https://depmap.org/portal/download/all. Cell lines were separated by lineage. Breast cancer cell lines were subcategorized by receptor status as in Suppl. Table 3.

### Survival analysis

An online tool for associating gene sets with breast cancer patient recurrence-free survival was used. Gene expression-based Outcome for Breast cancer Online (GOBO) (v.1.0.3) uses the datasets GSE1456, GSE2603, GSE6532, GSE3494, GSE7390, GSE12093, GSE2034, and GSE5327, and is available at https://co.bmc.lu.se/gobo. Both the Hallmark Oxidative Phosphorylation and KEGG Oxidative Phosphorylation gene sets were used for input. These gene sets are detailed in Suppl. Table 4.

### Statistical Testing

Cell growth data and IHC scores were analyzed by t-test (for 2-group experiments) or ANOVA followed by Bonferroni multiple comparison-adjusted post hoc testing between groups (for experiments with ≥3 groups). Tumor growth data were analyzed using the following linear mixed model on the logarithmic scale: Log_10_(tumor volume_it_) = a_i_ + b * t + e_it_, where i represents the i^th^ mouse, t represents time of tumor volume measurement, a_i_ represents the mouse-specific log tumor volume at t = 0, b represents the rate of tumor volume growth, and e_it_ represents deviation of measurements from the model over time^42,43^. Mouse heterogeneity (in baseline tumor volume) is represented by variance of a_i_, and b*log_e_(10) * 100 indicates tumor volume increase (%) per week. Treatment groups were compared using a z-test for slopes with standard error derived from the output of the function ‘lme’ from the library nlme in R. Synergy was determined as described in ref.^44^.

### Data availability

Sequencing data are available at NCBI SRA under accession PRJNA1023282 for WES and scRNA-seq. Proteomics data were deposited to ProteomeXchange under accession PXD048700 and to MassIVE under accession MSV0000093890. Metabolomics data were deposited to Metabolomics Workbench under accession ST003060.

## Results

### Endocrine therapy induces a persister state in ER+ breast cancer cells

Growth of cultured ER+ breast cancer cells is generally dependent upon estrogens present in serum (FBS). Hormone deprivation (HD) through culture in dextran/charcoal-stripped FBS (DCC-FBS) stunts the growth of estrogen-driven cells, but prolonged HD can lead to the emergence of proliferating long-term estrogen-deprived (LTED) derivatives as models of resistance to aromatase inhibitor-induced estrogen deprivation^45^. Modeling the acute effects of endocrine therapy to study persistence phenotypes, we subjected cells to (i) HD for up to 28 d, or (ii) treatment with the anti-estrogen fulvestrant for 7 d. ER+ cell lines displayed an initial period of growth followed by sustained stalling of growth that we define as “persistence” (Fig. 1A, Suppl. Fig. 1A/B). Accordingly, cells exposed to endocrine therapy displayed slower cell cycle progression (Suppl. Fig. 1C/D). This endocrine-tolerant persister state was reversible: when cells were re-exposed to the estrogen 17β-estradiol (E2) after 14 d of HD, growth resumed (Suppl. Fig. 1E); when a second cycle of HD was applied, growth again slowed. Indeed, the ER transcriptional response to E2 was maintained in endocrine-tolerant persisters (Suppl. Fig. 1F). Reseeding of cells after 0-28 d of HD showed that colony-forming capacity was slightly increased in persisters (Suppl. Fig. 1G), in agreement with our prior finding that persisters following estrogen deprivation in MCF7 xenografts were enriched for a stem-like phenotype^46,47^. Notably, the induction of endocrine-tolerant persistence did not alter sensitivity to conventional chemotherapeutics (Suppl. Fig. 1H), indicating that induction of endocrine tolerance does not activate broad drug persistence mechanisms.

**Figure 1:**
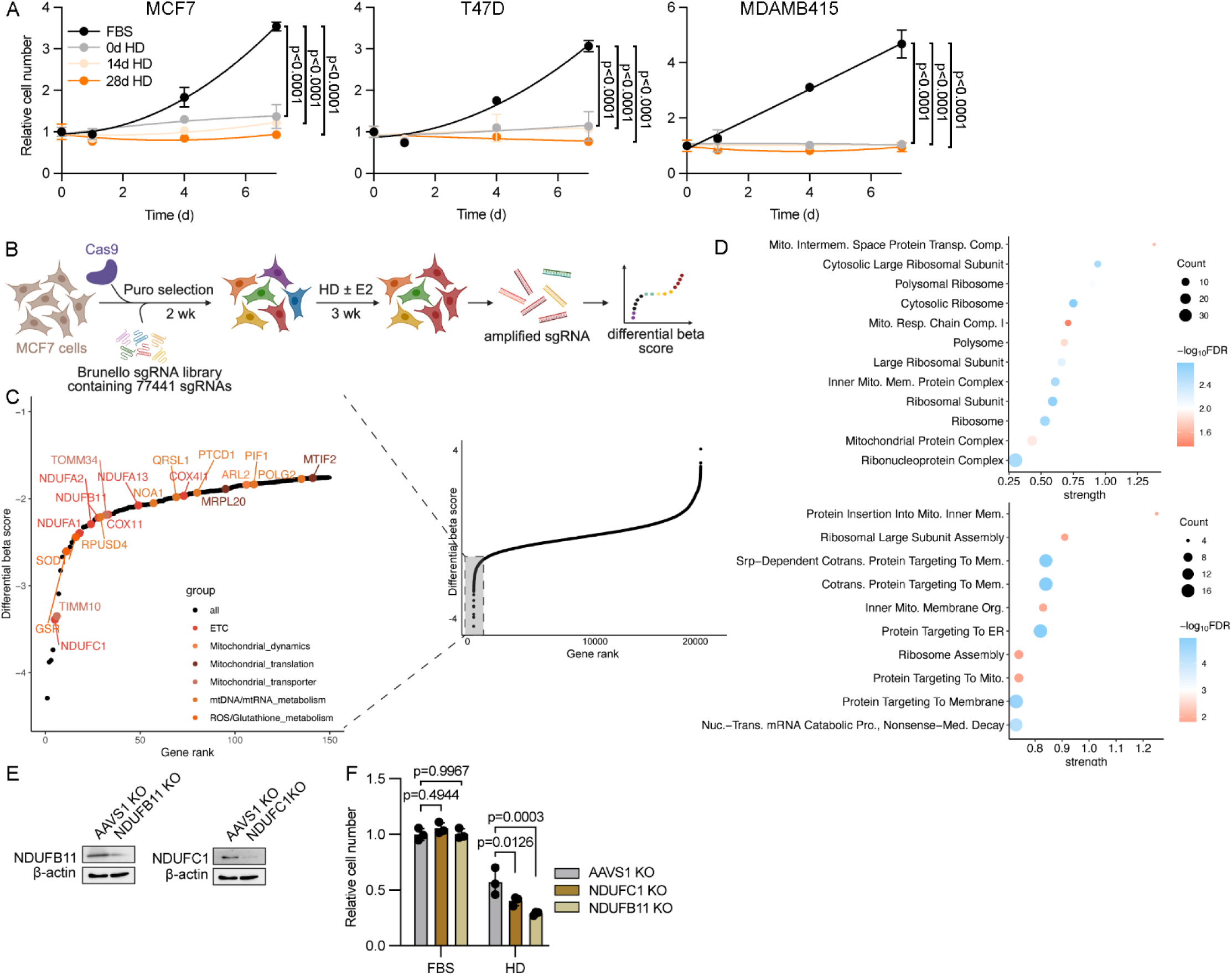
OXPHOS is a therapeutic vulnerability in endocrine-tolerant persisters. (A) Cells were pre-treated with growth medium (FBS) or hormone-depleted (HD) medium for up to 28 d, and then reseeded. Cells continued treatment, or were switched from growth to HD medium (“0 d HD”), and relative cell numbers were measured by serial imaging for 7 d. Curves were fit to a centered second-order polynomial. (B) Overview of nuclear genome-wide CRISPR knockout screening strategy. (C) Differential β-scores of all genes screened (*right*); inset at *left* shows genes with most negative scores. Genes with mitochondrial annotations are labeled with dots colored by function. (D) STRING enrichment analysis of the 500 most negatively enriched genes in HD cells from knockout screening for cellular components (*top*) and biological processes (*bottom*). (E) Immunoblot analysis of MCF7 cells with CRISPR knockout of *NDUFB11*, *NDUFC1*, or *AAVS1* (control). (F) Relative numbers of MCF7 cells cultured in growth or HD medium for 14 d. Data in (A) and (F) are shown as mean of 5-6 (A) or 3 (F) replicates ± SD, and were compared by one-way (F) or two-way (A) ANOVA with Bonferroni multiple comparison-adjusted posthoc test.

### Genome-wide screening reveals mitochondrial metabolism as a vulnerability in endocrine-tolerant persister cancer cells

To identify mechanisms of persistence to endocrine therapy, we conducted a genome-wide CRISPR knockout screen in MCF7 cells treated with HD ± E2 for 3 wk (Fig. 1B). Differential β-scores (β_HD_ – β_HD+E2_) were calculated to identify genes essential for persistence (Fig. 1C, Suppl. Table 5). Enrichment analysis of the top 500 genes negatively selected in HD conditions highlighted mitochondrial pathways (Fig. 1D). Genes encoding proteins associated with electron transport chain complex I such as *NDUFC1* and *NDUFB11* had highly negative differential β-scores (Fig. 1C). ER+ breast cancer cell lines showed higher dependency on *NDUFC1* and *NDUFB11* than most other cancer types and other breast cancer subtypes (Suppl. Fig. 2A), suggesting that mitochondrial metabolism is more essential in ER+ breast cancer. Validation experiments showed that knockout of *NDUFC1* or *NDUFB11* decreased mitochondrial membrane polarization and spare respiratory capacity compared to control cells (Fig 1E, Suppl. Fig. 2B-D), suggesting that knockout of a single complex I subunit does not disrupt basal respiration but only maximal respiratory capacity. NDUFC1 or NDUFB11 loss conferred negative selection in HD but not E2-replete conditions (Fig. 1F), suggesting that complex I becomes a vulnerability during persistence.

### ER+ breast cancer persister cells do not show enrichment for genetic alterations and arise stochastically

To evaluate the landscape of clonal composition during endocrine therapy, we explored how genetic alterations arise in an MCF7 xenograft model. We previously demonstrated that E2-driven MCF7 tumors regress upon E2 withdrawal in ovariectomized NOD/SCID/Il2γR^−/−^ (NSG) mice and yield a stable slow-cycling persister cell population after 60-90 d^47^. If maintained for ≥1 yr, a fraction of mice can develop E2-independent recurrent tumors. We harvested tumors and residual tumor beds from ovariectomized mice (i) during E2-driven growth (baseline), (ii) after 90 d of E2 deprivation (persistent state), and (iii) after recurrence during E2 deprivation (resistant state) (Fig. 2A). Tumor DNA was analyzed by whole exome sequencing (WES) to profile genomic alterations. Compared to baseline tumors, we found few copy number alterations (CNAs) in persistent tumor cells, but CNAs were more abundant in resistant tumors (Fig. 2B, Suppl. Fig. 2A, Supp. Table 6). We detected 1,295 single nucleotide variants (SNVs) across 11 tumors (Suppl. Fig. 2B, Supp. Table 7). Focusing on variability in variant allele frequency between tumors, we found that like the architecture of CNAs, differences in variants arose between resistant tumors and baseline/persistent tumors (Fig. 2C).

**Figure 2:**
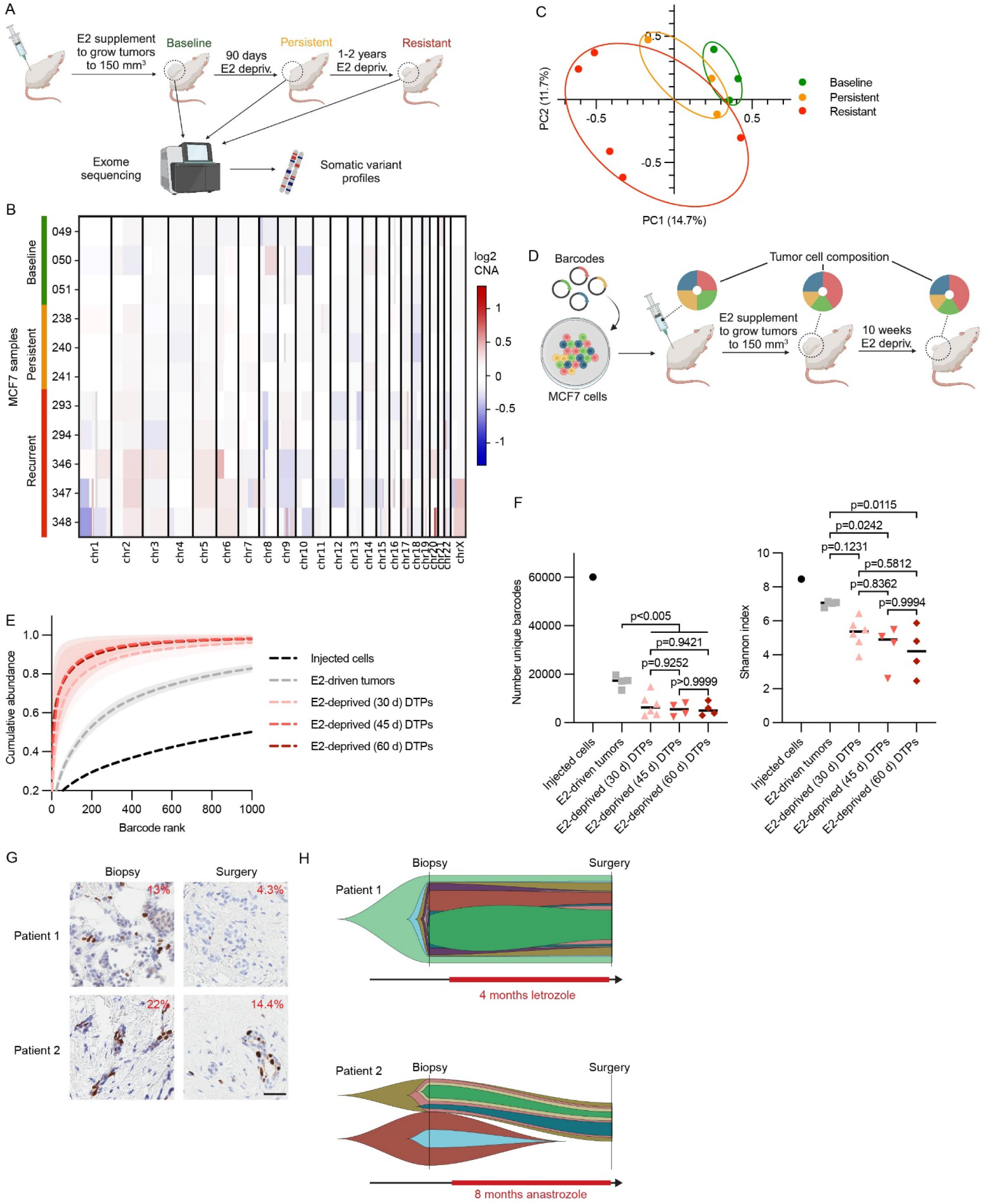
Persistence during endocrine therapy occurs stochastically and is not driven by genetic predisposition. (A) Overview of growth, treatment, harvest, and WES of MCF7 tumors. (B) Heatmap of CNAs of MCF7 tumors. (C) Principal component analysis of SNV profiles. (D) Overview of experimental strategy for clonal analysis of MCF7 xenografts over time. (E) Cumulative abundance of the 1,000 most abundant barcodes in each treatment group. Shaded regions represent 95% CIs. (F) Numbers of unique barcodes detected (*left*) and Shannon diversity index of barcodes (*right*) were compared by one-way ANOVA with Bonferroni multiple comparison-adjusted posthoc test. Horizontal bars indicate medians. (G) Representative images showing Ki67 IHC of breast tumor specimens obtained from patients before and after 4 or 8 months of neoadjuvant therapy with an aromatase inhibitor. Proportions of positively stained cells are indicated therein. (H) Fish plots of variant clusters determined by PyClone and ordered by ClonEvol derived from WES of tumor samples in (G).

In addition to WES, we used an orthogonal approach of genetic barcoding to temporally track clonal composition. Barcoded MCF7 cells were orthotopically implanted into ovariectomized mice, and tumor growth was induced with exogenous E2 (Fig. 2D). Nineteen whole tumors were harvested from mice at baseline and after 30-60 d of E2 deprivation (persistent state). Barcode sequencing revealed different clonal subsets overrepresented in different tumors, indicating that if clonal selection occurred, then it was stochastic (Suppl. Fig. 2C). Barcode diversity progressively decreased during E2 deprivation, reaching full cumulative abundance with fewer barcodes (Fig. 2E). Indeed, numbers of unique barcodes and Shannon diversity index temporally decreased (Fig. 2F), suggesting that E2 deprivation imposed selection on cells. However, we observed high similarity in barcode abundance profiles across tumors and time points, supporting the notion that overall clonal composition remains stable during treatment (Suppl. Fig. 2D).

We extended the strategy of inferring clonal composition to clinical samples from two patients with early-stage ER+/HER2-breast cancer. We performed WES on tumor specimens obtained at diagnosis (baseline biopsy) and surgery following 4 or 8 months of neoadjuvant aromatase inhibitor therapy (Supp. Tables 8-9). Ki67 immunohistochemistry (IHC) on tumor sections confirmed that endocrine treatment decreased malignant cell proliferation (Fig. 2G). SNVs were used to infer clonal population structures (Fig. 2H, Suppl. Fig. 2E). Patient 1’s tumor exhibited limited clonal changes during treatment. Patient 2’s tumor contained two clones that were extinguished during treatment, while proportions of other clones remained stable. These data collectively suggest that genetic alterations do not broadly drive selection of endocrine-tolerant persisters, but endocrine therapy may remove a subset of clones.

### Endocrine-tolerant persisters shift metabolic programming towards mitochondrial oxidative phosphorylation (OXPHOS)

To explore the functional context of persister metabolism, we quantitatively analyzed the proteomes of MCF7 cells treated with HD ± E2 for 4 wk, or HD for 3 wk followed by E2 treatment for 1 wk to revert cells from persistence. Proteomic profiles of E2-reverted cells were distinct from those of cells treated with continuous E2, possibly due to acute effects of ER reactivation (Suppl. Fig. 4A/B). Gene set coregulation analysis^48^ indicated that the Hallmark Estrogen Response Early and Late gene sets exhibit high correlation scores, indicating a strong response of ER activity downregulation from HD (Suppl. Fig. 4C), validating our approach to using quantitative proteomics to capture biological state.

Proteomic profiling revealed decreased glycolytic protein levels in HD persisters, but levels of proteins associated with mitochondria including complex I and mitochondrial ribosomal subunits were maintained (Fig. 3A). HD persisters showed increased mitochondrial content and polarization that were reversed upon restoration of E2 (Fig. 3B/C, Suppl. Fig. 4D-F). Pretreatment with fulvestrant elicited similar effects to HD (Fig. 3D), suggesting that metabolic reprogramming is driven by loss of ER activity. Measurement of the levels of representative electron transport chain proteins showed a substantial HD-induced increase in the complex IV component COXII that is encoded by mitochondrial DNA (Fig. 3E), suggesting increased abundance of mitochondria.

**Figure 3:**
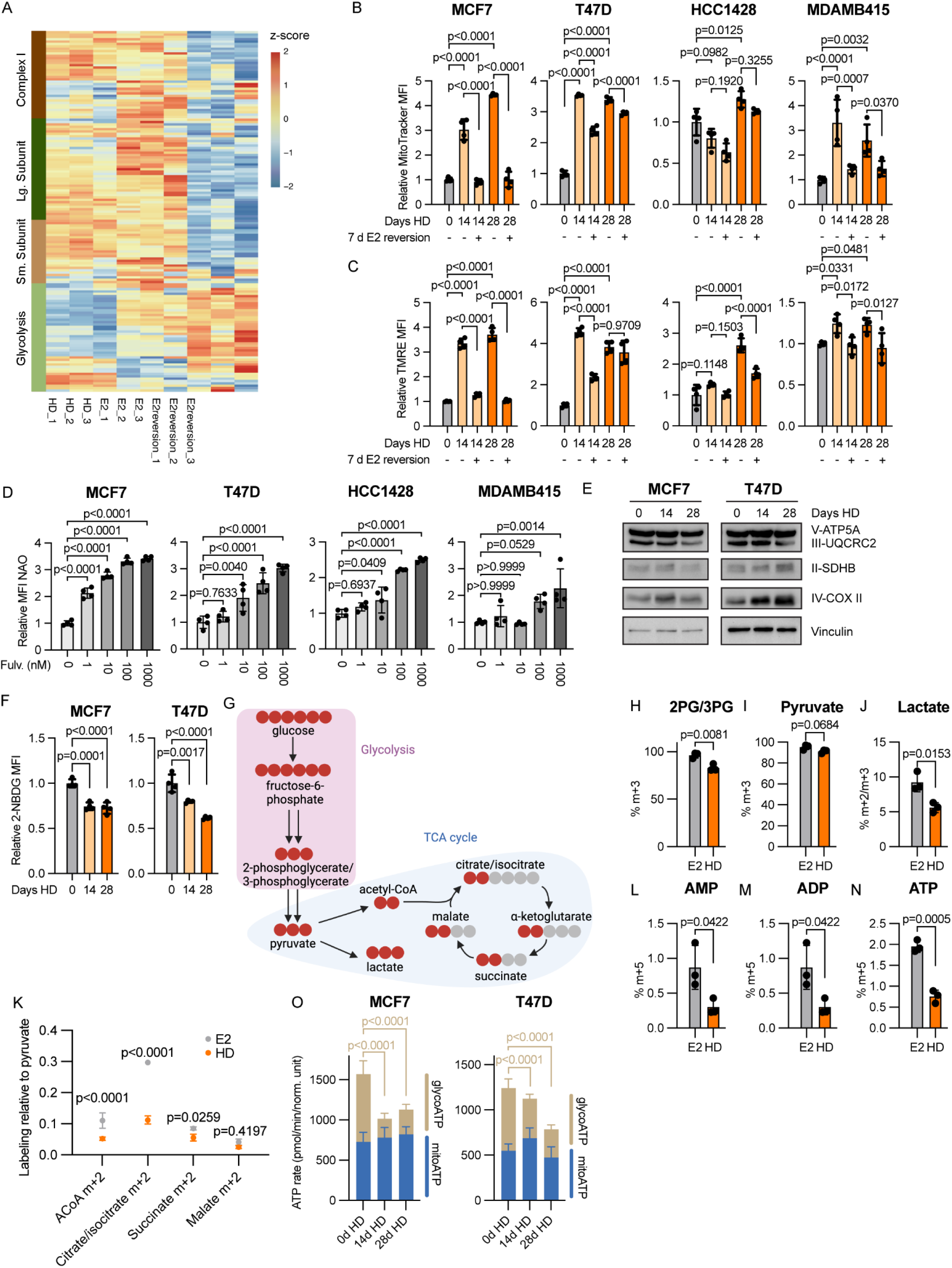
Endocrine-tolerant persisters undergo metabolic reprogramming and rely on mitochondria for ATP production. (A) Quantitative proteomic analysis of MCF7 cells treated with HD ± 1 nM E2 for 28 d, or with HD for 21 d followed by restoration of E2 for 7 d. The heatmap shows relative levels of proteins stratified by complex I, mitochondrial ribosomes, or glycolysis. (B/C) Flow cytometry analysis of cells pre-treated with HD for 0-28 d, and then ± E2 for 7 d. Cells stained with MitoTracker Deep Red (B) or TMRE (C) were assayed by flow cytometry, and median fluorescence intensities are shown. (D) Flow cytometry analysis of cells treated with 0-1 μM fulvestrant for 7 d, stained with NAO, and analyzed as in (B). (E) Immunoblot analysis of cells treated with HD for 0-28 d. (F) Flow cytometry analysis of cells treated with HD for 0-28 d, labeled with 2-NBDG, and analyzed as in (B). (G) Overview of carbon tracing using [U-^13^C]-glucose with mass spectrometry. Red circles denote ^13^C. (H-J/L-N) Metabolite abundances of MCF7 cells treated with HD for 3 wk followed by HD ± 1 nM E2 for 1 wk. Cells were labeled with [U-^13^C]-glucose for 45 min. Metabolites were extracted and quantified by mass spectrometry. (K) Metabolite abundances normalized to pyruvate abundances. (O) Cells were treated with HD for 0-28 d prior to Seahorse assay to measure ATP generation rate from mitochondria or glycolysis (*n* = 14-16/group). Data were analyzed by t-test (H-N) or one-way ANOVA with Bonferroni multiple comparison-adjusted posthoc test (B-D/F/O). Data are presented as mean ± SD.

We tested whether the endocrine therapy-induced increases in mitochondrial content and decreases in glycolytic protein levels (Fig. 3A/B) corresponded with decreased glycolytic function. Indeed, HD persisters showed decreased glucose uptake compared to non-HD controls (Fig. 3F). Since OXPHOS is dependent upon intermediate metabolites produced by glycolysis and the tricarboxylic acid (TCA) cycle, we postulated that alterations in metabolite availability limit the contribution of glycolysis to sustaining endocrine-tolerant persisters. We conducted ^13^C-glucose tracing to measure intermediate metabolite flux in cells treated with HD for 3 wk, followed by treatment ± E2 for 1 wk, and labeled for 45 min (Fig. 3G, Suppl. Table 10). Metabolite abundances associated with glycolysis and the TCA cycle were decreased in HD persisters (Fig. 3H-K). Interestingly, metabolites downstream of pyruvate in the TCA cycle exhibited less labeling in HD cells, suggesting metabolic bottlenecks at multiple steps (Fig. 3K). This limited glycolytic flux in persisters was reflected in decreased adenine nucleotide production (Fig. 3L-N). Furthermore, ATP generated from glycolysis fell in persisters, setting the stage for mitochondrial respiration as the principal source of ATP (Fig. 3O, Suppl. Fig. 4G). We postulated that high mitochondrial content could predispose cells to becoming persisters. Surprisingly, both mitochondria^high^ and mitochondria^low^ cells exhibited similar endocrine sensitivity (Suppl. Fig. 4H/I), suggesting that metabolic reprogramming in endocrine-tolerant persisters is an induced/acquired phenotype.

### Higher mitochondrial content is associated greater residual proliferative capacity in human breast tumors

We conducted clinical trial NCT04568616 to determine whether residual cancer cells following neoadjuvant endocrine therapy exhibit alterations in markers of mitochondria and metabolism compared to baseline ER+/HER2-breast tumors. Thirty-two evaluable patients received neoadjuvant therapy with the aromatase inhibitor letrozole for 2-16 wk (Fig. 4A, Suppl. Fig. 5A). Tumor tissue specimens collected before (diagnostic biopsy) and after (surgical specimen) letrozole treatment were analyzed by IHC for Ki67, the mitochondrial marker MT-CO2, and the fatty acid oxidation enzyme CPT1α. Neoadjuvant endocrine therapy decreased Ki67 score in 31/32 cases (Suppl. Fig. 5B). Higher Ki67 scores after short-term endocrine therapy predict earlier recurrence and worse long-term disease outcome^49,50^. Higher post-letrozole Ki67 scores were significantly correlated with letrozole-induced increases in MT-CO2 (Fig. 5B/C) but not CPT1α (Suppl. Fig. 5C), suggesting that tumors with higher likelihood of recurrence acquired increases in mitochondrial content during the course of endocrine therapy. Notably, MT-CO2 histoscore did not correlate with Ki67 score across all samples (Suppl. Fig. 5D), indicating that mitochondrial content is not a general correlate of proliferative state. Analysis of an integrated dataset of primary tumor transcriptomic profiles from 914 breast cancer patients^51^ further revealed that transcriptional signatures of OXPHOS were prognostic of shorter recurrence-free survival (Suppl. Fig. 5E/F).

**Figure 4:**
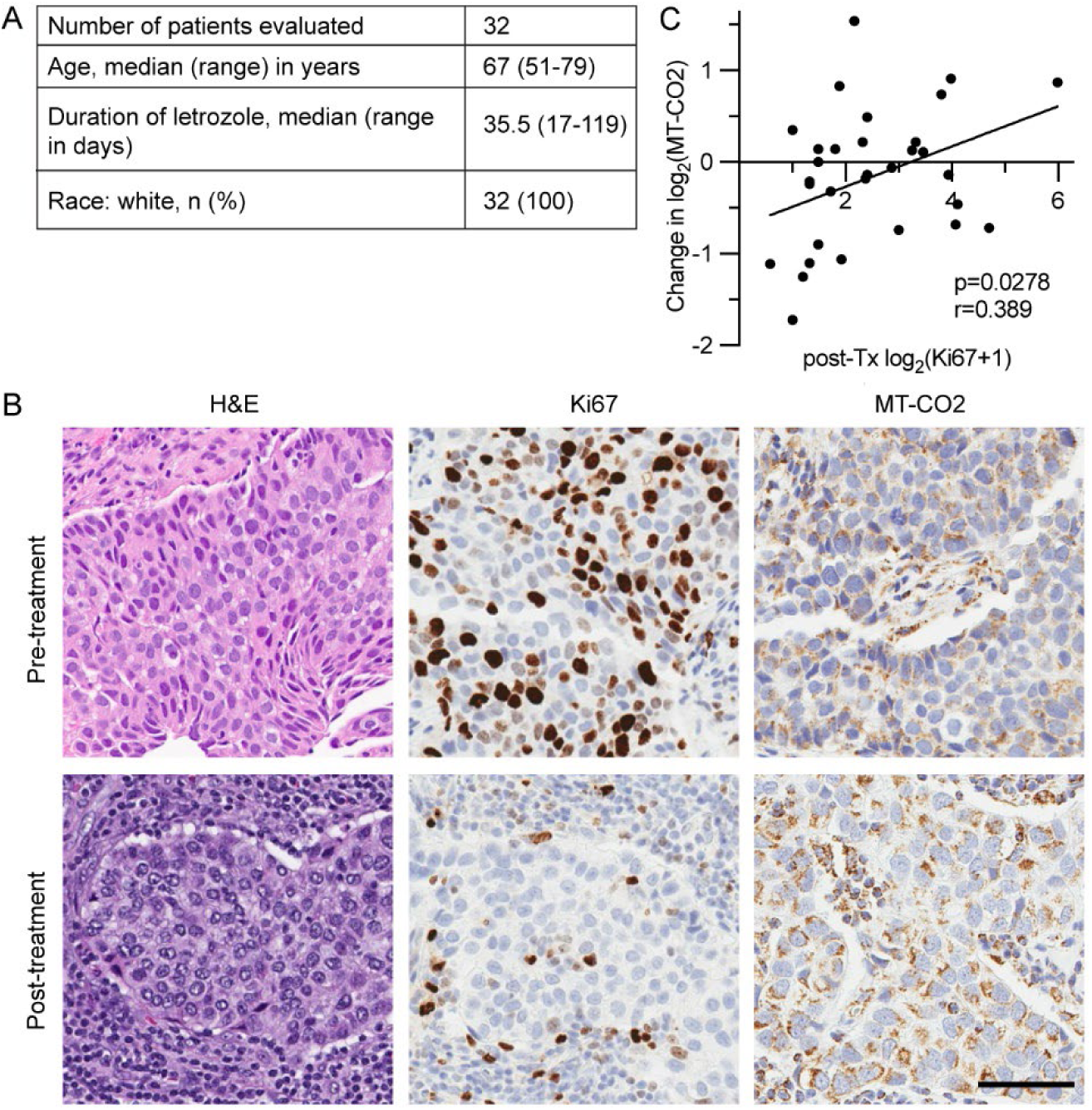
Mitochondrial content is upregulated during endocrine therapy in primary ER+/HER2-breast tumors that exhibit residual cancer cell proliferation. (A) Patient cohort characteristics. (B) Representative images of patient-matched pre-and post-letrozole breast tumor specimens stained with H&E and IHC for Ki67 and MT-CO2. Scale bar is 100 μm. (C) Plot of changes in MT-CO2 score (post-letrozole minus pre-letrozole) versus post-treatment Ki67 score analyzed by Pearson correlation.

**Figure 5:**
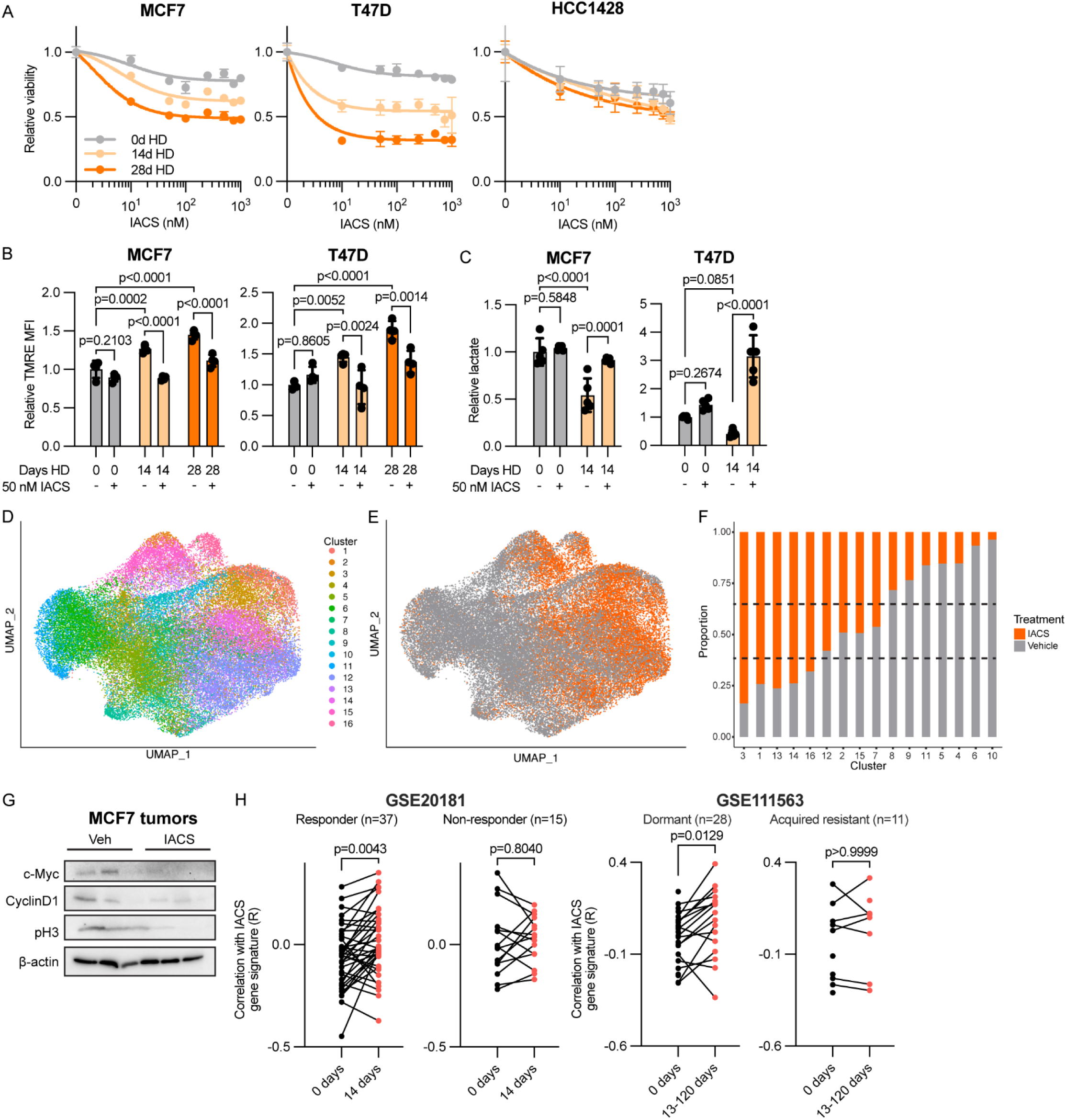
Inhibition of mitochondrial OXPHOS selectively targets endocrine-tolerant persisters. (A) Quantification of viability of cells pre-treated with HD for 0-28 d, then reseeded and treated with HD and a dose range of IACS-010759. Relative cell numbers were measured 7 d later (*n* = 6/group). (B) Flow cytometry analysis of cells pre-treated with HD for 0-28 d, and then treated ± IACS for 3 d before labeling with TMRE. (C) Quantification of excreted lactate normalized to relative cell number as determined by SRB assay. Cells were treated as in (B). (D) Uniform manifold and projection (UMAP) of cells subjected to scRNA-seq. Ovariectomized mice were orthotopically implanted with MCF7/miRFP720 cells and a s.c. E2 pellet. When tumors reached 200 mm^3^, E2 pellets were removed. After 60 d of E2 deprivation, mice were treated ± IACS for 3 d. Tumors (*n*=2/group) were harvested and dissociated, and ∼15,000 cancer cells/tumor were isolated by flow cytometry for scRNA-seq. Colors indicate *k*-means clusters. (E) UMAP projection by treatment condition of cells from (D). (F) Breakdown by cluster of results from (D/E). Dotted horizontal lines indicate thresholds used to determine whether a cluster was predominated by a treatment group. (G) Immunoblot analysis of E2-deprived MCF7 tumors from mice treated ± IACS for 50 d. (H) A 123-gene signature was derived from MCF-7 tumor analyzed in (F) to reflect transcriptional changes induced by IACS. This signature was compared with transcriptional profiles of human tumors serially sampled during neoadjuvant endocrine therapy in two patient cohorts stratified by response. Pearson correlation R values were compared by Bonferroni multiple comparison-adjusted posthoc test. In (A-C), data are presented as mean ± SD.

### Inhibition of mitochondrial complex I selectively targets endocrine-tolerant persisters

Since endocrine-tolerant persisters showed upregulation of mitochondrial content (Figs. 3B/D and 4C), and an increased proportion of ATP was being derived from respiration (Fig. 3O), we tested dependence upon OXPHOS using compounds targeting mitochondrial function. Tigecycline is an FDA-approved anti-bacterial drug, but in eukaryotic cells tigecycline inhibits mitochondrial ribosomes^52^. Tigecycline more effectively inhibited growth of endocrine-tolerant persisters than parental controls (Suppl. Fig. 6A).

Since CRISPR screening implicated complex I in persistence (Fig. 1), we tested IACS-010759 (IACS) as a selective electron transfer inhibitor of complex I recently evaluated in clinically^53–55^. IACS more effectively suppressed growth of HD persisters than parental controls in 2/3 cell lines (Fig. 5A). IACS reduced mitochondrial respiration and returned polarization to the baseline levels seen in parental cells (Fig. 5B, Suppl. Fig. 6B), suggesting that mitochondrial alterations induced by endocrine therapy contribute to IACS sensitization.

Inhibition of OXPHOS can force cells to utilize glycolysis as a bioenergetic pathway^56^. HD persisters were unable to maintain their elevated mitochondrial polarization under IACS (Fig. 5B), likely causing a metabolic switch towards glycolysis as reflected by IACS-induced increases in lactate production (Fig. 5C). Furthermore, this switch results in lower ATP levels in persisters when treated with IACS as opposed to controls (Suppl. Fig. 6C), suggesting that IACS provoked an energetic crisis that suppressed growth (Fig. 5A). Blockade of both OXPHOS (with IACS) and glycolysis (with galactose) further decreased persister viability (Suppl. Fig. 6D).

Since complex I initiates the electron transport chain, and inhibition by IACS can hinder the function of subsequent complexes, we sought to discern the necessity of complex I function for endocrine-tolerant persistence. We generated T47D cells stably expressing *Saccharomyces cerevisiae* NADH-quinone oxidoreductase (NDI1), which can replace the human complex I function of electron transport from NADH to ubiquinone without proton translocation^57^. NDI1 increased cellular oxygen consumption rate (OCR) and lowered extracellular acidification rate (ECAR), suggesting boosted mitochondrial function (Suppl. Fig. 6E). NDI1 partially rescued endocrine-tolerant persisters from the growth-suppressive effects of IACS (Suppl. Fig. 6F), suggesting that complex I function drives persistence.

We sought to infer the metabolic state of endocrine-tolerant persisters under IACS treatment in human breast tumors through transcriptomic profiling. To first generate a transcriptomic signature reflective of the metabolic shift induced by complex I inhibition, mice bearing E2-induced MCF7 tumors were treated with estrogen deprivation for 60 d to yield persisters^47^, and then treated ± IACS for 3 d. Tumors (n=2/group) were dissociated, and ∼15,000 cancer cells/tumor were isolated for single-cell RNA sequencing (scRNA-seq). Uniform Manifold Approximation and Projection (UMAP) revealed 16 major clusters of cells (Fig. 5D), some of which were dominated by one treatment group (Fig. 5E/F, Suppl. Fig. 6G), suggesting the emergence of a new transcriptional state in response to IACS.

Gene set enrichment analysis of the combined clusters 1, 3, 13, 14, and 16 (composed mostly of IACS-treated cells) compared to all other clusters revealed gene sets related to metabolism and cell cycle (Suppl. Table 11). Querying for differential gene expression between two groups of clusters predominated by one treatment (clusters 1, 3, 13, 14, 16 vs. 4, 5, 6, 8, 9, 10,11) revealed 123 transcripts (Suppl. Table 12). Only 34 transcripts were upregulated in IACS-treated cells including *SLC39A6*, *MLPH*, *C6orf141*, *DST*, and *TNFRSF11B* (Suppl. Fig. 6H), which are associated with less aggressive breast cancer and correlated with improved survival^58–62^. IACS treatment decreased expression of proliferative drivers (*MYC* and *CCND1*) and the complex I genes *NDUFB9* and *NDUFS5* (Suppl. Fig. 6I). We validated that IACS decreased protein levels of c-Myc, Cyclin D1, and phospho-histone H3_S10_ (proliferative marker) in MCF7 tumors (Fig. 5G) and T47D cells (Suppl. Fig. 6J), indicating that IACS further arrests cell cycling beyond the effects of estrogen deprivation.

Since genes induced by IACS have been associated with favorable prognosis (Suppl. Fig. 6H), we evaluated the correlation between transcriptional changes induced by IACS in MCF7 tumor persisters (Fig. 6E/F) and transcriptional profiles of human tumors sampled during neoadjuvant endocrine therapy in cohorts GSE20181 and GSE111563^39,40^. Tumor specimens used for transcriptomic profiling were obtained at baseline and during aromatase inhibitor treatment. Patients were stratified based on clinical response to treatment determined by imaging (Responder/Dormant vs. Non-Responder/Acquired Resistance). Correlations of the IACS-induced signature and tumor gene expression profiles increased during aromatase inhibitor treatment in cases that did not presurgically develop endocrine resistance (i.e., Responders/Dormant), suggesting that IACS induces a less aggressive phenotype (Fig. 5G). Exclusion of genes related to cell cycle did not alter these associations (Suppl. Fig. 6K). Together, these data suggest an effective strategy of selectively targeting endocrine-tolerant persister cells through complex I inhibition.

**Figure 6:**
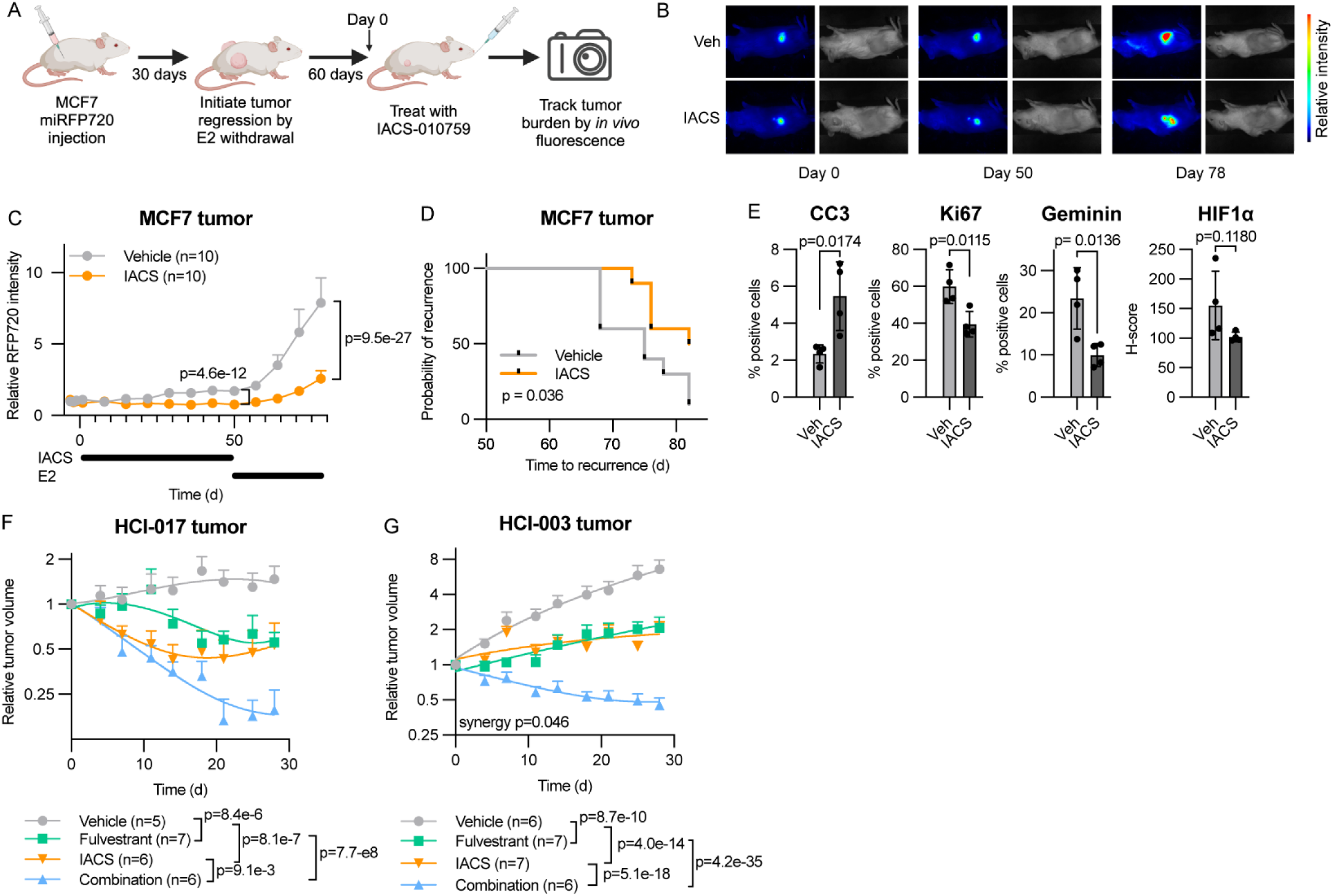
OXPHOS inhibition enhances endocrine therapy in ER+ breast cancer xenografts. (A) Overview of MCF7/miRFP720 study of IACS efficacy. Ovariectomized mice were orthotopically implanted with MCF7/miRFP720 cells and a s.c. E2 pellet. When tumors reached 200 mm^3^, E2 was withdrawn for 60 d. Mice were then randomized to treatment with vehicle or IACS starting on “Day 0” for 50 d. Relative tumor burden was measured by fluorescence imaging. A new E2 pellet was implanted on Day 51, and mice were monitored for recurrence (defined as tumors regrowing to 50 mm^3^). (B) Representative serial mouse-matched images of miRFP720 fluorescence and white light. (C) Quantification of relative miRFP720 fluorescent signal serially measured in mice treated as in (A). (D) Kaplan-Meier curves of tumor recurrence compared by log-rank test. (E) IHC analysis of tumors harvested after 50 d IACS treatment. Data were analyzed by one-way ANOVA with Bonferroni multiple comparison-adjusted posthoc test. (F-G) Quantification of tumor volume in mice bearing orthotopic HCI-003 and HCI-017 PDX tumors that were randomized to treatment with vehicle, fulvestrant, IACS, or the combination when tumors reached 150 mm^3^. Data in (C/F/G) are shown as mean ± SEM and were analyzed by linear mixed modeling.

### Complex I inhibition decreases tumor burden during and following endocrine therapy

Since luciferase activity is dependent upon ATP availability^63^, we instead utilized miRFP720 as a near-infrared red fluorescent protein reporter^64^ to non-invasively monitor tumor burden *in vivo*. Ovariectomized mice with orthotopically implanted MCF7/miRFP720 cells were treated with exogenous E2 to induce tumor growth, and then E2 was withdrawn for 60 d to yield endocrine-tolerant persisters (Fig. 6A). Persisters displayed a high fluorescent signal-to-background ratio (Suppl. Fig. 7A), and serial measurements across 3 d showed low inter-day variability (Suppl. Fig. 7B). Mice bearing MCF7/miRFP720 endocrine-tolerant persisters were randomized to treatment ± IACS for 50 d. Serial measurements showed that IACS significantly decreased persister cell burden compared to vehicle-treated control (Fig. 6B/C). Following treatment ± IACS, mice were again challenged with exogenous E2. Mice that had received IACS showed lower tumor regrowth rate during E2 re-challenge (Fig. 6C/D, Suppl. Fig. 7C/D), indicating that IACS treatment of mice with endocrine-tolerant persisters decreased tumor-reforming potential. IACS increased apoptosis (cleaved caspase 3), decreased proliferation (Ki67, geminin), and decreased levels of hypoxia-inducible factor-1α (HIF-1α, marker of OXPHOS inhibition^65^) in tumor persisters (Fig. 6E, Suppl. Fig. 7E).

Since persister cell burden (i.e., residual disease) cannot be serially measured *in vivo* without genetic engineering, we also tested the effects of complex I inhibition in the context of concurrent endocrine therapy in translationally relevant endocrine-sensitive patient-derived xenograft (PDX) models (HCI-017, HCI-003). Single-agent treatment with fulvestrant or IACS stunted tumor growth, and the combination elicited the greatest reduction in tumor volume (Fig. 6F/G, Suppl. Fig. 8A/B), supporting the notion that complex I inhibition is more effective in metabolically reprogrammed persister cells. Endpoint molecular analysis of tumors after 28 d of treatment showed that fulvestrant and combination treatments reduced proliferation (Ki67, geminin) in residual HCI-017 tumors, and the combination did so in HCI-003 tumors, but drug-induced apoptosis was not evident at this late time point (Suppl. Fig. 8C-F). These findings support the strategy of targeting mitochondrial OXPHOS in combination with endocrine therapy to decrease the burden of endocrine-tolerant persisters and prevent tumor recurrence in ER+ breast cancer.

## Discussion

Despite the widespread use of adjuvant endocrine therapy, a significant proportion of patients with ER+ breast cancer experience recurrence. Thus, there is a critical need to develop improved treatment strategies that prevent recurrence. The years-to-decades-long latency between (A) initial diagnosis and treatment (e.g., surgery, radiotherapy) and (B) clinically evident recurrence of ER+ breast cancer indicates that a subpopulation of clinically occult cancer cells persists that can ultimately give rise to recurrence despite endocrine therapy. Herein, we describe a subpopulation of endocrine-tolerant persister cancer cells that undergo metabolic reprogramming to rely on mitochondrial OXPHOS as the main source of ATP to support viability. These cells appear to be stochastically selected for persistence without differences in mitochondrial content or genetic predisposition; such genetic findings are in agreement with prior reports on other cancer types^5^. Endocrine-tolerant persisters are sensitized to OXPHOS inhibition, which reduced tumor regrowth potential. Endocrine therapy with fulvestrant synergized with the OXPHOS inhibitor IACS-010759 to induce regression of ER+ PDX tumors, indicating that endocrine therapy and OXPHOS inhibition are most effective in combination.

Endocrine-tolerant persisters showed upregulation of mitochondrial content and polarization (Figs. 3B/C), increased proportion of ATP generation from OXPHOS (Fig. 3O), sensitivity to mitochondrial complex I inhibition (Figs. 5A and 7C/F/H), and decreased glycolysis (Fig. 3A/F/O) in preclinical models. Mitochondrial content was also upregulated during neoadjuvant endocrine therapy in human ER+/HER2-breast tumors with poor prognosis (i.e., high residual Ki67^49,50^) (Fig. 4C). Upregulation and dependency of persister cells on mitochondrial metabolism has been observed in other cancer types^66–70^, suggesting a commonality of cellular metabolic reprogramming that may yield a broadly applicable therapeutic strategy to eradicate persisters. Such reprogramming is reflective of a shift from the proliferative “Warburg-like” state of high glycolylic flux to a more “differentiated-like” state of OXPHOS to produce ATP^71^.

We observed that inhibition of mitochondrial complex I decreased the tumor regrowth potential of endocrine-tolerant persisters in estrogen-deprived mice (Fig. 6C/D) as a model of residual disease following (neo)adjuvant aromatase inhibitor therapy in addition to potentiating fulvestrant in PDX models (Fig. 6F/G). El-Botty et al. recently described dose-dependent responses to IACS in subcutaneous PDX tumors derived from patients’ ER+ breast cancer bone metastases, and showed that the combination of fulvestrant and the CDK4/6 inhibitor palbociclib (approved for metastatic disease) enhanced response to IACS^72^. Together, these findings suggest that mitochondrial metabolism may be therapeutically tractable in both endocrine-sensitive and endocrine-resistant disease settings, each in the context of an endocrine therapy backbone.

A key question in the cancer field is whether primary tumors can provide biomarkers that predict mechanisms of persistence and resistance, which would inform treatment decisions. In ER+ breast cancer, high residual tumor cell proliferation following neoadjuvant endocrine therapy is predictive of earlier recurrence^49,50^ and has been considered for recommendation of additional systemic therapy^73^. We found transcriptional signatures reflective of high OXPHOS in treatment-naïve primary ER+ breast tumors to be associated with worse outcome (Suppl. Fig. 5E/F). These data suggest that early molecular signs of impending endocrine persistence that portend poor disease outcome are detectable in primary tumors and may be useful as a biomarker to guide early intervention to target persisters. In addition, El-Botty et al. observed that PI3K pathway-activating mutations in *PIK3CA* or *AKT1* are associated with sensitivity to IACS^72^. Consistent with this, the lone PI3K pathway-wild-type model tested herein (HCC-1428 cells) was less sensitive to IACS compared to other cell lines (MCF-7, T47D) and PDX models (HCI-003, HCI-017) that all bore *PIK3CA* activating mutations (Figs. 5A and 6C/F/G). In addition to the well-described role of PI3K pathway activation in endocrine resistance^45^, these observations suggest that (i) PI3K pathway activation may drive the metabolic reprogramming observed under endocrine therapy, and (ii) such mutations may offer genetic biomarkers to predict sensitivity to OXPHOS inhibitors and guide designs for neoadjuvant basket trials.

One limitation of this study was the characterization of metabolic reprogramming being limited to *in vitro* models. Technical developments are needed to enable comprehensive *in vivo* assessment of functional metabolic parameters (e.g., flux), which will enable the evaluation of pharmacodynamic effects of drugs on metabolism in tumor and non-tumor tissues. Drugs targeting mitochondrial metabolism continue to be clinically evaluated as anti-cancer agents, including devimistat as an inhibitor of the TCA cycle enzymes pyruvate dehydrogenase and alpha-ketoglutarate dehydrogenase. Since metabolic pathways are often shared between cancer and non-cancer cells, identifying strategies to increase therapeutic index *in vivo* are critical to the development of metabolism-targeted therapeutics. A strategy being developed that has shown increased therapeutic index in animal studies is conjugating OXPHOS inhibitors with triphenylphosphonium cations to drive accumulation in mitochondria^74^. Given the emerging pan-cancer theme of metabolic reprogramming towards mitochondrial metabolism as a feature of persister cells, continued development of tolerable therapeutic strategies is warranted.

## Acknowledgements

This work was supported by NIH (R01CA200994, R01CA267691, R01CA262232, and R01CA211869 to TWM; Rosalind Borison Memorial Fund and F31CA278418 to ST; F31CA220936 to RAH; the Center for Quantitative Biology at Dartmouth P20GM130454; Dartmouth College Cancer Center Support Grant P30CA023108; S10OD030242). We thank the patients and their families for participating in this research, and the following Dartmouth Cancer Center Shared Resources for their support: Mouse Modeling; Pathology (RRID: SCR_023479); Biostatistics; Bioinformatics; Microscopy; Genomics & Molecular Biology (RRID: SCR_021293); Office of Clinical Research. Metabolomics measurements were performed by the Michigan Regional Comprehensive Metabolomics Resource Core.

