## Supplementary material for "Endocrine persistence in ER+ breast cancer is accompanied by metabolic vulnerability in oxidative phosphorylation": Suppl Figs 1-6

**SUPPLEMENTAL DATA**

**Supplemental Figure 1: ER+ breast cancer cells persist under endocrine therapy.** (A) Relative numbers of cells plated in quadruplicate and treated with growth medium (FBS) or hormone-depleted (HD) medium and assayed by serial imaging. Curves were fit to a centered second-order polynomial, and t-test of coefficients was performed. (B) Relative numbers of cells plated in quadruplicate in growth medium treated +/- fulvestrant for 14 d, followed by fulvestrant withdrawal for 14 d. $^{\#}p<0.0001$ comparing cells treated with fulvestrant (all doses) vs. control during Days 0-14 by ANOVA with Bonferroni multiple comparison adjustment. For cells pre-treated with fulvestrant, ANOVA with Bonferroni multiple comparison-adjusted posthoc test p-values are shown for Day 28 vs. 14. (C/D) Cell cycle profiling was performed after 0-28 d of HD (C), or after 7 d of treatment +/- 1 uM fulvestrant (D). Proportions of cells in S-phase were compared by ANOVA with Bonferroni multiple comparison-adjusted posthoc test (C) or t-test (D). (E) Quantification of relative numbers of cells measured at each time point (inverted black triangles in schematic). Cells were treated with HD for 14 d (pink line), re-seeded and treated with 1 nM E2 (gray line), and re-seeded and treated with HD for 14 d (red line). p-values were calculated by ANOVA with Bonferroni multiple comparison adjustment of relative cell numbers after 7 d of growth ($n$ = 5-6/group). (F) Quantification of luciferase activity of cells pre-treated with HD for 0-28 d, and then transfected with an ERE-firefly luciferase reporter plasmid and a CMV-*Renilla* control plasmid. Cells were treated +/- 1 nM E2 for 1 d, and luciferase activities were measured. Firefly luciferase signal was normalized to *Renilla* signal, and data are shown as relative luminescence units (RLU). Groups were compared by t-test. (G) Quantification of cells pre-treated with HD for 0-28 d, and then re-seeded in HD medium with 1 nM E2. Relative numbers of cells were measured 14 d later and analyzed by Cytation 5 image analysis. Representative images are shown. (H) Quantification of relative viability of cells pre-treated with HD for 0-28 d, and then reseeded. Cells were treated with dose ranges of drugs as indicated for 7 d, and relative cell numbers were measured.

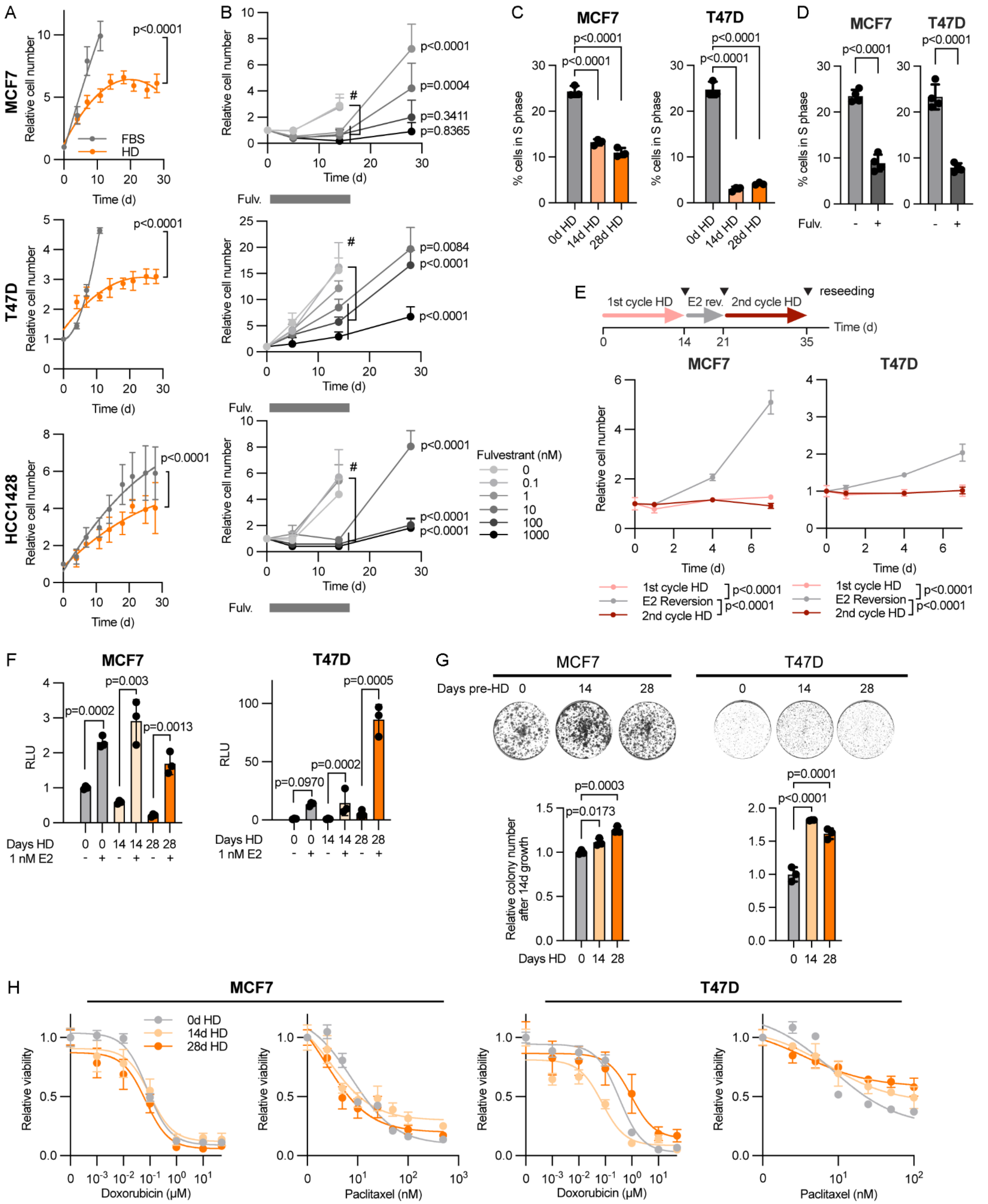
A
MCF7
Relative cell number
p<0.0001
FBS
HD
Time (d)
T47D
HCC1428
B
Relative cell number
#
p<0.0001
p=0.0004
p=0.3411
p=0.8365
Fulv.
p=0.0084
p<0.0001
Fulvestrant (nM)
0
0.1
1
10
100
1000
C
% cells in S phase
0d HD
14d HD
28d HD
D
Fulv.
E
1st cycle HD
E2 rev.
2nd cycle HD
reseeding
E2 Reversion
F
RLU
p=0.0002
p=0.003
p=0.0013
p=0.0970
p=0.0005
Days HD
1 nM E2
G
Days pre-HD
p=0.0003
p=0.0173
Relative colony number after 14d growth
p=0.0001
H
Relative viability
Doxorubicin (μM)
Paclitaxel (nM)

**Supplemental Figure 2: Pan-cancer genome-wide CRISPR screening identifies *NDUFC1* and *NDUFB11* as vulnerabilities in ER+ breast cancer.** (A) Gene effect scores extracted from DepMap project of cell lines categorized by cancer type. (B) Flow cytometry analysis of cells in MCF7 cells in which CRISPR was used to knock out NDUFC1, NDUFB11, or AAVS1 (control). Cells were stained with TMRE, and signal was measured by flow cytometry. Relative median fluorescence intensities (MFI) are shown. (C) Serial measurements of oxygen consumption rate (OCR) in cell lines from (B) as analyzed by Seahorse MitoStress test kit (n = 20-22/group). (D) Components of oxygen consumption as calculated from data shown in (C). Data in (B/D) are shown as mean ± SD and were compared by ANOVA with Bonferroni multiple comparison-adjusted posthoc test.

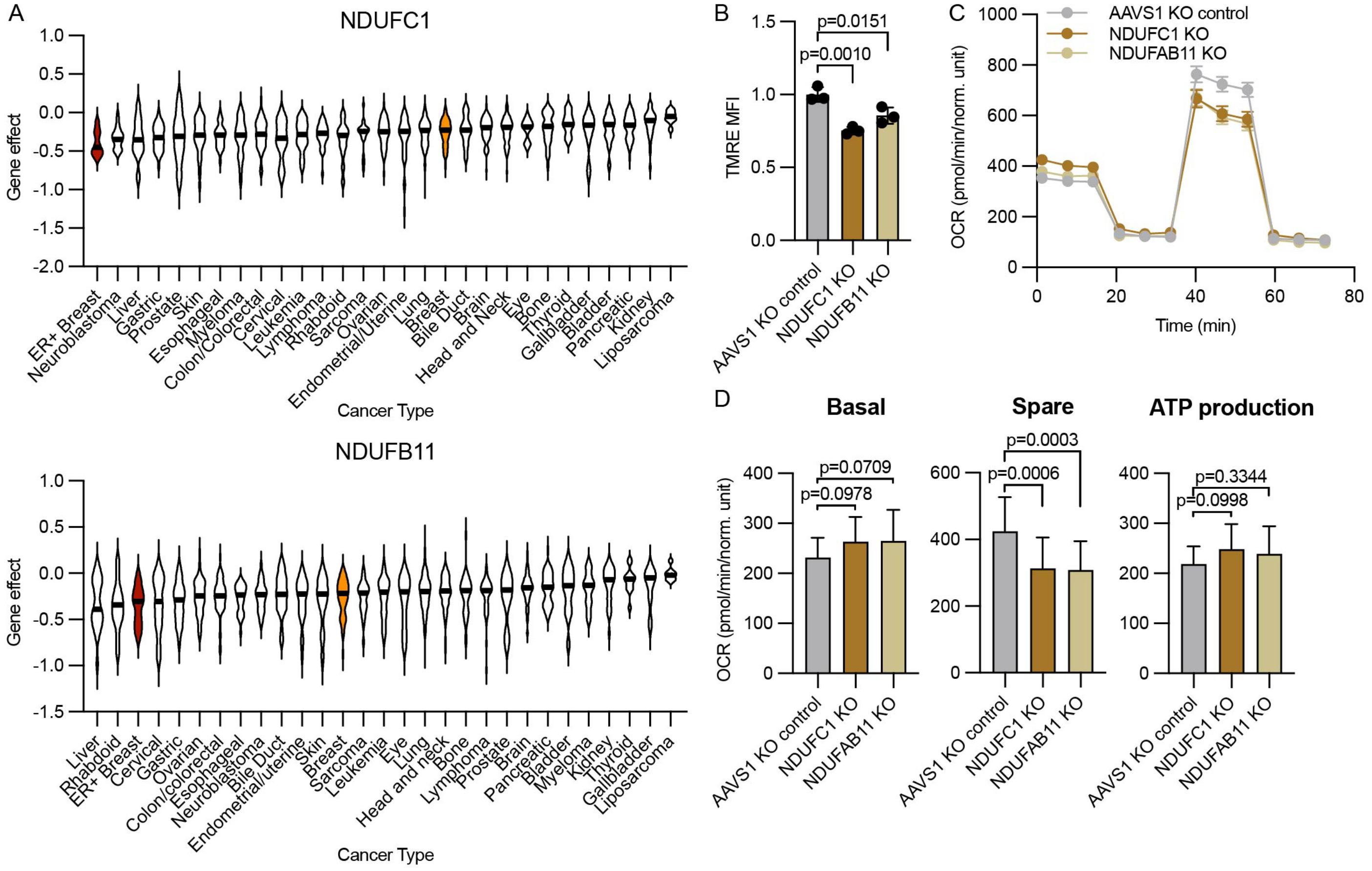

**Supplemental Figure 3: Ontogeny of endocrine-tolerant persistence.**

(A) PCA plot of CNAs in MCF7 xenografts harvested before, during, and following recurrence on E2 deprivation and analyzed by WES. (B) Heatmap of SNV VAFs detected in tumors from (A). (C) Heatmap showing barcode abundance in MCF7 cells genetically barcoded and orthotopically implanted with a s.c. E2 pellet. When tumors reached 200 $mm^3$, E2 was withdrawn. Tumors were harvested after 0-60 of E2 deprivation, and barcodes were sequenced and quantified. Abundances are depicted as row-wise z-scores. (D) Pearson correlation heatmap of barcode abundances across tumors from (C). (E) VAF of variants grouped by PyClone analysis of WES data from breast tumors specimens obtained from two patients before and after 4 or 8 months of neoadjuvant therapy with an aromatase inhibitor.

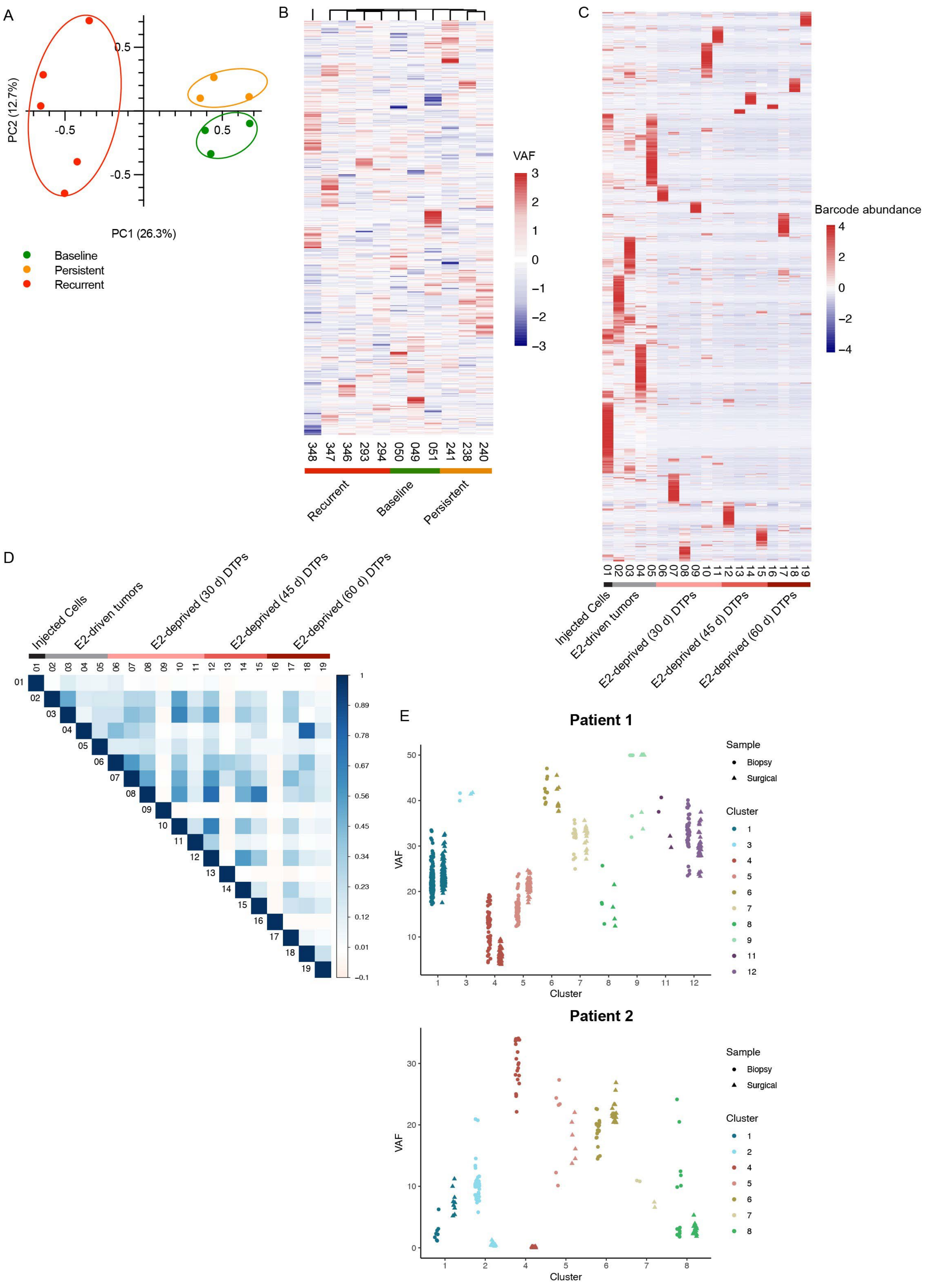
A
PC2 (12.7%)
PC1 (26.3%)
0.5
-0.5
Baseline
Persistent
Recurrent
B
VAF
348 347 346 293 294 050 049 051 241 238 240
Recurrent
Baseline
Persisrtent
C
Barcode abundance
01 02 03 04 05 06 07 08 09 10 11 12 13 14 15 16 17 18 19
Injected Cells
E2-driven tumors
E2-deprived (30 d) DTPs
E2-deprived (45 d) DTPs
E2-deprived (60 d) DTPs
D
Injected Cells
E2-driven tumors
E2-deprived (30 d) DTPs
E2-deprived (45 d) DTPs
E2-deprived (60 d) DTPs
1
0.89
0.78
0.67
0.56
0.45
0.34
0.23
0.12
0.01
−0.1
E
Patient 1
VAF
Cluster
Sample
Biopsy
Surgical
Cluster
1 3 4 5 6 7 8 9 11 12
Patient 2
VAF
Cluster
Sample
Biopsy
Surgical
Cluster
1 2 4 5 6 7 8

**Supplemental Figure 4: Endocrine-tolerant persisters exhibit increased mitochondrial content and polarization.** (A) Quantitative proteomic analysis of MCF7 cells treated with HD +/- 1 nM E2 for 28 d, or with HD for 21 d followed by restoration of E2 for 7 d. The heatmap shows relative levels of proteins with ≥2 matching peptides. (B) PCA plot of protein abundance across all proteins depicted in (A). (C) Gene set co-regulation analysis of Hallmarks pathways using protein abundance data from (A). Red arrows highlight Estrogen Early and Late Response gene sets. (D) Flow cytometry analysis of cells pre-treated with HD for 0-28 d, and then +/- E2 for 7 d. Cells stained with NAO were assayed by flow cytometry, and median fluorescence intensities (MFI) were calculated. (E) Flow cytometry analysis of cells treated with HD for 0-28 d, and then stained with anti-Tom20-AlexaFluor594 conjugated antibody followed by analysis as in (D). (F) Flow cytometry analysis of cells treated with HD for 0-28 d, stained with JC-1, and analyzed by flow cytometry to measure ratios of polarized (red) to depolarized (green) mitochondria across the population of cells. (G) Serial oxygen consumption rate (OCR) measurements of cells treated with HD for 0-28 d, reseeded, and analyzed by Seahorse MitoStress test kit. Components of oxygen consumption were calculated ($n$ = 15-16/group). Data in (D-G) are shown as mean ± SD and were compared by ANOVA with Bonferroni multiple comparison-adjusted posthoc test. (H) Overview of flow cytometry sorting strategy for cells with high or low mitochondrial content by MitoTracker staining intensity. (I) Viability analysis of cells following sorting as in (H), re-seeded, and treated with a dose range of fulvestrant for 7 d.

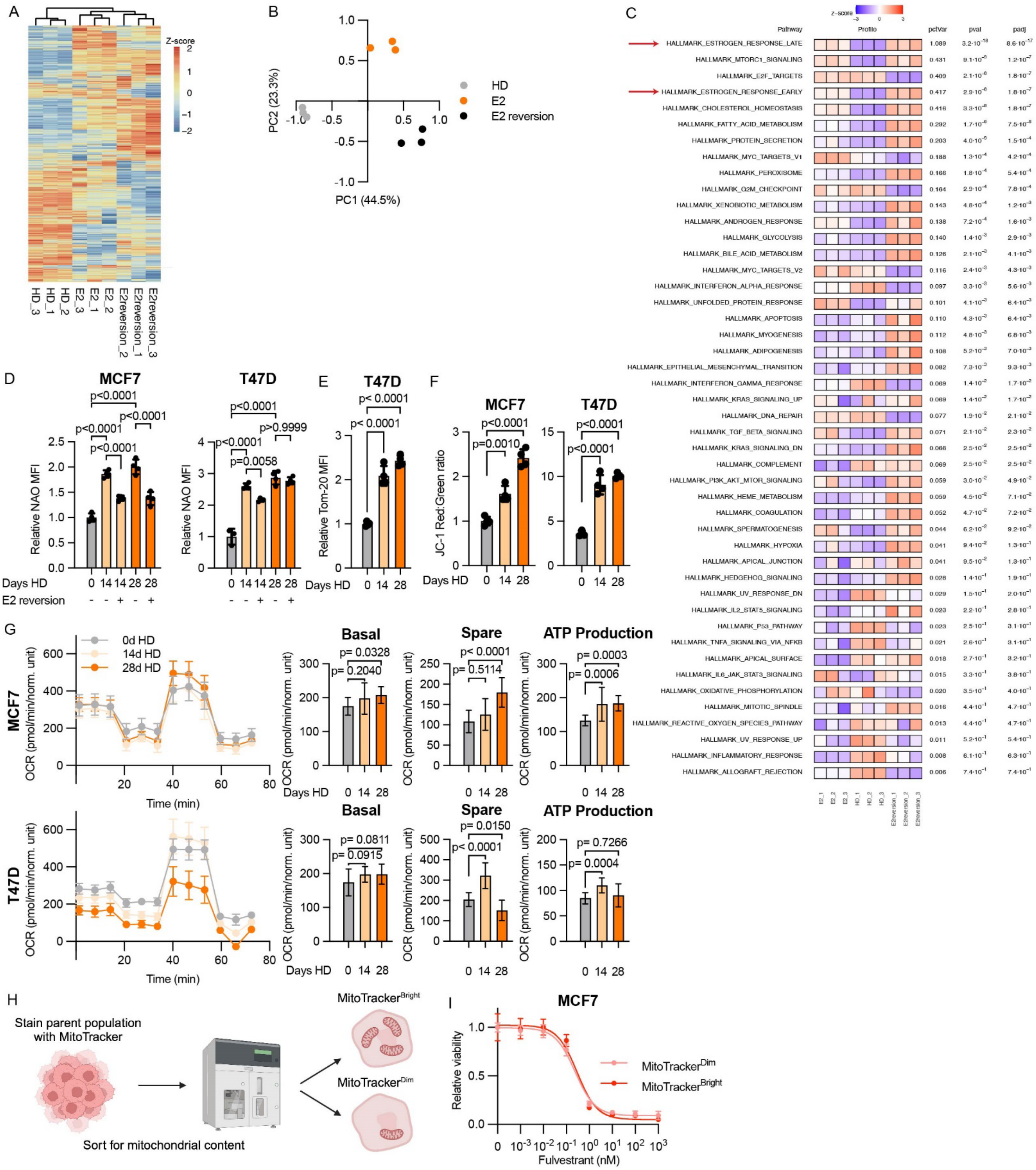
A
Z-score
HD_3
HD_1
HD_2
E2_3
E2_1
E2_2
E2reversion_2
E2reversion_1
E2reversion_3
B
PC2 (23.3%)
PC1 (44.5%)
HD
E2
E2 reversion
C
Pathway
Profile
pctVar
pval
padj
HALLMARK_ESTROGEN_RESPONSE_LATE
HALLMARK_MTORC1_SIGNALING
HALLMARK_E2F_TARGETS
HALLMARK_ESTROGEN_RESPONSE_EARLY
HALLMARK_CHOLESTEROL_HOMEOSTASIS
HALLMARK_FATTY_ACID_METABOLISM
HALLMARK_PROTEIN_SECRETION
HALLMARK_MYC_TARGETS_V1
HALLMARK_PEROXISOME
HALLMARK_G2M_CHECKPOINT
HALLMARK_XENOBIOTIC_METABOLISM
HALLMARK_ANDROGEN_RESPONSE
HALLMARK_GLYCOLYSIS
HALLMARK_BILE_ACID_METABOLISM
HALLMARK_MYC_TARGETS_V2
HALLMARK_INTERFERON_ALPHA_RESPONSE
HALLMARK_UNFOLDED_PROTEIN_RESPONSE
HALLMARK_APOPTOSIS
HALLMARK_MYOGENESIS
HALLMARK_ADIPOGENESIS
HALLMARK_EPITHELIAL_MESENCHYMAL_TRANSITION
HALLMARK_INTERFERON_GAMMA_RESPONSE
HALLMARK_KRAS_SIGNALING_UP
HALLMARK_DNA_REPAIR
HALLMARK_TGF_BETA_SIGNALING
HALLMARK_KRAS_SIGNALING_DN
HALLMARK_COMPLEMENT
HALLMARK_PI3K_AKT_MTOR_SIGNALING
HALLMARK_HEME_METABOLISM
HALLMARK_COAGULATION
HALLMARK_SPERMATOGENESIS
HALLMARK_HYPOXIA
HALLMARK_APICAL_JUNCTION
HALLMARK_HEDGEHOG_SIGNALING
HALLMARK_UV_RESPONSE_DN
HALLMARK_IL2_STAT5_SIGNALING
HALLMARK_P53_PATHWAY
HALLMARK_TNFA_SIGNALING_VIA_NFKB
HALLMARK_APICAL_SURFACE
HALLMARK_IL6_JAK_STAT3_SIGNALING
HALLMARK_OXIDATIVE_PHOSPHORYLATION
HALLMARK_MITOTIC_SPINDLE
HALLMARK_REACTIVE_OXYGEN_SPECIES_PATHWAY
HALLMARK_UV_RESPONSE_UP
HALLMARK_INFLAMMATORY_RESPONSE
HALLMARK_ALLOGRAFT_REJECTION
D
MCF7
T47D
Relative NAO MFI
Days HD
E2 reversion
E
T47D
Relative Tom-20 MFI
Days HD
F
MCF7
T47D
JC-1 Red:Green ratio
Days HD
G
0d HD
14d HD
28d HD
MCF7
T47D
OCR (pmol/min/norm. unit)
Time (min)
Basal
Spare
ATP Production
Days HD
H
Stain parent population with MitoTracker
Sort for mitochondrial content
MitoTracker^Bright
MitoTracker^Dim
I
MCF7
Relative viability
Fulvestrant (nM)
MitoTracker^Dim
MitoTracker^Bright

**Supplemental Figure 5: Patients with primary human ER+/HER2- breast tumors with a transcriptional signature of high OXPHOS have worse outcomes.** (A) CONSORT flow diagram of NCT04568616 clinical trial patients. (B) Ki67 scores of patients' tumor specimens acquired pre- and post-neoadjuvant letrozole. (C) Plot of changes in CPT1$\alpha$ score (post-letrozole minus pre-letrozole) versus post-treatment Ki67 scores analyzed by Pearson correlation. (D) IHC scores for Ki67 and MT-CO2 of all pre- and post-letrozole breast tumor specimens analyzed by Pearson correlation. (E/F) Kaplan-Meier curves of recurrence-free survival (RFS) in a multi-study patient cohort with any subtype (*left*) or ER+ subtype (*right*) stratified by median average expression of Hallmark Oxidative Phosphorylation (C) and KEGG Oxidative Phosphorylation (D) gene sets. Curves were compared by log-rank test.

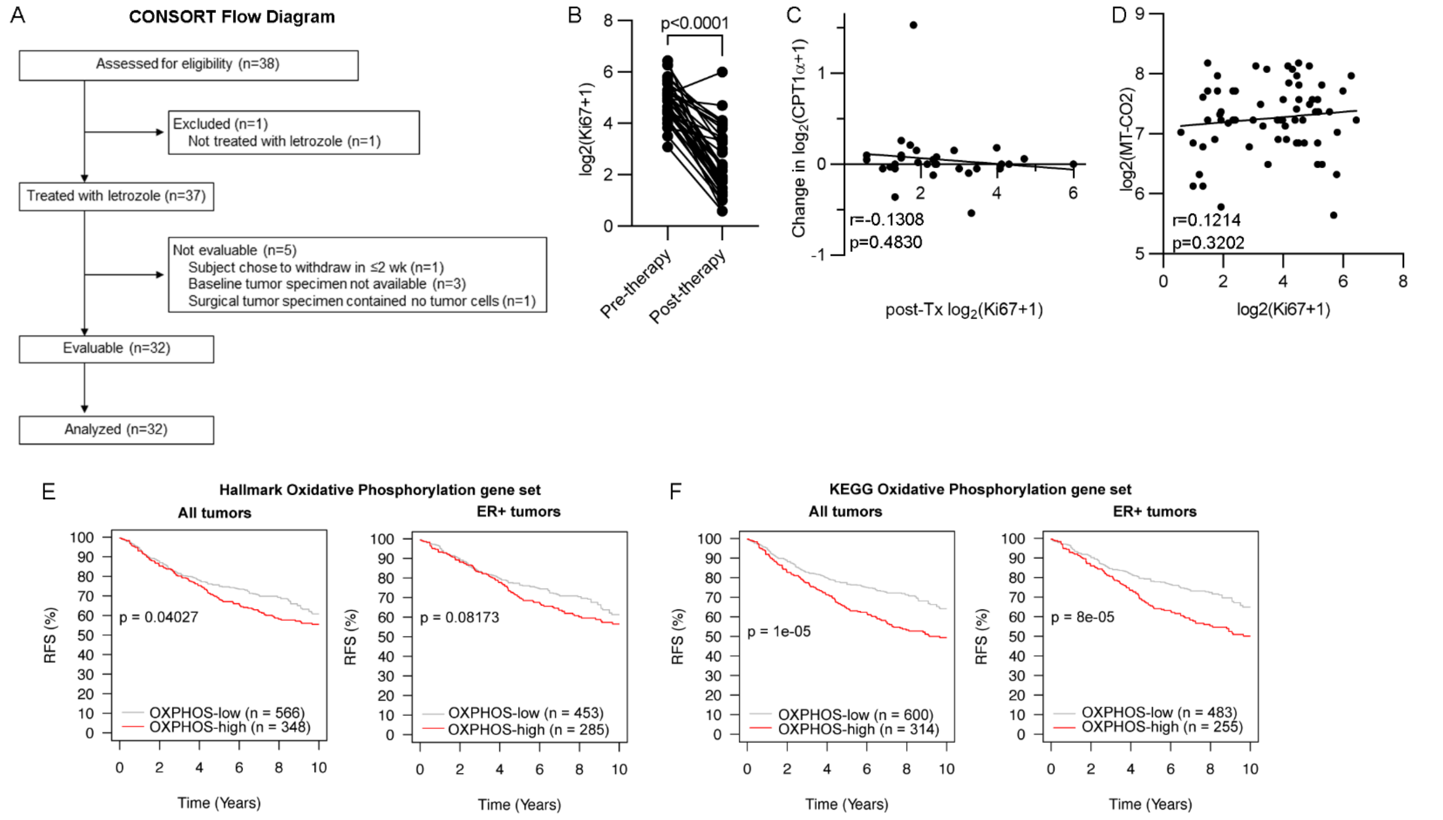

**Supplemental Figure 6: NDI1 promotes resistance to OXPHOS inhibition.** (A) Relative viability of cells pre-treated with HD for 0 or 14 d, and then reseeded and treated with a dose range of tigecycline for 10 d in HD medium. Relative numbers of cells were measured after 7 d of drug exposure (*n* = 6/group). (B) OCR of cells pre-treated with HD for 0 or 14 d, and then treated +/- IACS for 1 d. Cells were analyzed by Seahorse to measure OCR. (C) Quantification of ATP concentration in cells pre-treated with HD for 0-28 d, and then treated +/- IACS for 3 d. Relative ATP levels were measured by CellTiter-Glo luminescence assay, and luciferase units were normalized to relative cell numbers (determined by SRB in wells with matched conditions). (D) Relative numbers of cells treated with HD medium containing either 4.5 g/L glucose or galactose, and/or IACS-010759 for 14 d. (E) OCR of T47D cells stably expressing NDI1 or vector control were analyzed as in (B) to measure OCR and ECAR. (F) Quantification of relative numbers of cells from (E) pre-treated with either 14 d of HD (*left*) or 7 d of 1 uM fulvestrant (*right*). Cells were then reseeded and treated ± 50 nM IACS-010759 for 7 d. (G) Proportions of cells from each treatment group (± IACS) within each *k*-means cluster are shown from scRNA-seq. Ovariectomized mice were orthotopically implanted with MCF7/miRFP720 cells and a s.c. E2 pellet. When tumors reached 200 $mm^3$, E2 pellets were removed. After 60 d of E2 deprivation, mice were treated +/- IACS for 3 d. Tumors were harvested and dissociated, and ~15,000 cancer cells/tumor were isolated by FACS for scRNA-seq. (H/I) Gene-level expression data extracted from scRNA-seq data. (J) Immunoblot analysis of cells pre-treated with HD for 0-28 d, and then treated +/- IACS for 3 d. (K) Nine genes with a Gene Ontology annotation associated with the cell cycle were removed from the 123-gene transcriptional signature induced by IACS in estrogen-deprived MCF7 tumors. This cell cycle-depleted 114-gene signature (in Suppl. Table 12) was compared with transcriptional profiles of human tumors serially sampled during neoadjuvant endocrine therapy in two patient cohorts stratified by response. Pearson correlation R values were compared by mixed-modeling followed by Bonferroni multiple comparison-adjusted posthoc test. In (A-F), data are presented as mean ± SD. Data were analyzed by ANOVA with Bonferroni multiple comparison-adjusted posthoc test (B/D), t-test (C/E/F/K), and Wilcoxon signed-rank test (H/I).

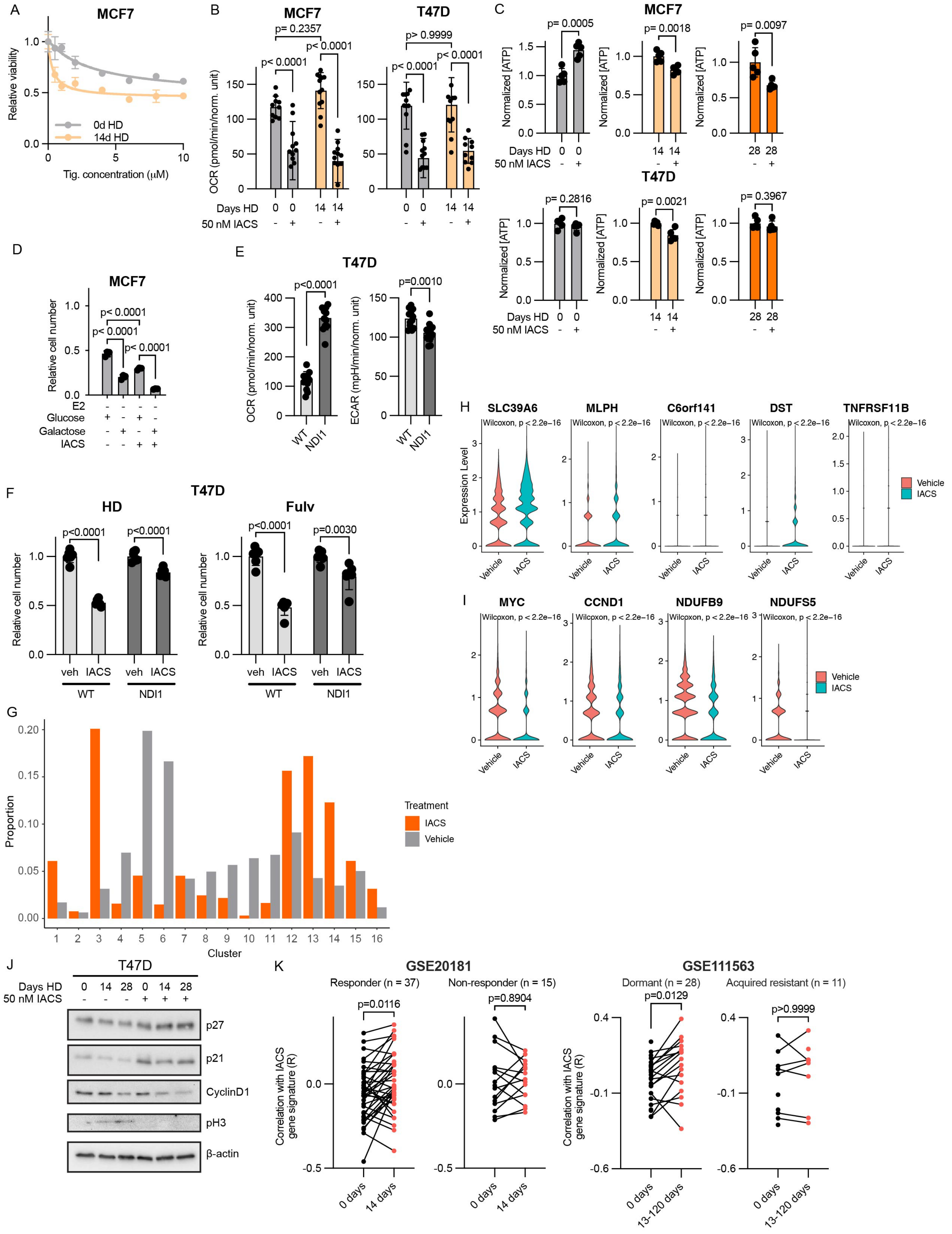
A
MCF7
Relative viability
0d HD
14d HD
Tig. concentration (μM)
B
MCF7
T47D
p= 0.2357
p< 0.0001
p< 0.0001
p< 0.0001
p> 0.9999
p< 0.0001
p< 0.0001
OCR (pmol/min/norm. unit)
Days HD
50 nM IACS
C
MCF7
p= 0.0005
p= 0.0018
p= 0.0097
Normalized [ATP]
T47D
p= 0.2816
p= 0.0021
p= 0.3967
D
MCF7
p< 0.0001
p< 0.0001
p< 0.0001
Relative cell number
E2
Glucose
Galactose
IACS
E
T47D
p<0.0001
p=0.0010
OCR (pmol/min/norm. unit)
ECAR (mpH/min/norm. unit)
WT
NDI1
F
T47D
HD
Fulv
p<0.0001
p<0.0001
p<0.0001
p=0.0030
Relative cell number
veh
IACS
WT
NDI1
G
Proportion
Cluster
Treatment
IACS
Vehicle
H
SLC39A6
MLPH
C6orf141
DST
TNFRSF11B
Wilcoxon, p < 2.2e−16
Expression Level
Vehicle
IACS
I
MYC
CCND1
NDUFB9
NDUFS5
Wilcoxon, p < 2.2e−16
Vehicle
IACS
J
T47D
Days HD
50 nM IACS
p27
p21
CyclinD1
pH3
β-actin
K
GSE20181
Responder (n = 37)
Non-responder (n = 15)
p=0.0116
p=0.8904
Correlation with IACS gene signature (R)
0 days
14 days
GSE111563
Dormant (n = 28)
Acquired resistant (n = 11)
p=0.0129
p>0.9999
13-120 days

**Supplemental Figure 7: OXPHOS inhibition synergizes with endocrine therapy in MCF7 breast tumors.** (A) Plot depicting signal from a mouse bearing MCF7/miRFP720 orthotopic xenografts evaluated by fluorescence imaging. A representative intensity plot of miRFP720 signal detected in regions of tumor, whole mouse, and the machine stage (around the mouse) shows specificity of fluorescent signal from tumor. (B) Fluorescence signal acquired over 3 consecutive d from 20 tumors that had been HD for 60 d. Data are shown as mean of 3 data points ± SD. (C) Relative burden of individual MCF7/miRFP720 tumors from fluorescence imaging. “Day 0” reflects the start of treatment +/- IACS after 60 d of E2 deprivation. (D) Tumor volume of MCF7/miRFP720 xenografts induced to regrow by implantation of a new s.c. E2 pellet. Data are shown as mean ± SEM and were analyzed by linear mixed modeling. (E) Representative IHC images of MCF7/miRFP720 tumors harvested at the end of IACS-010759 treatment (Day 50). Scale bar = 50 um.

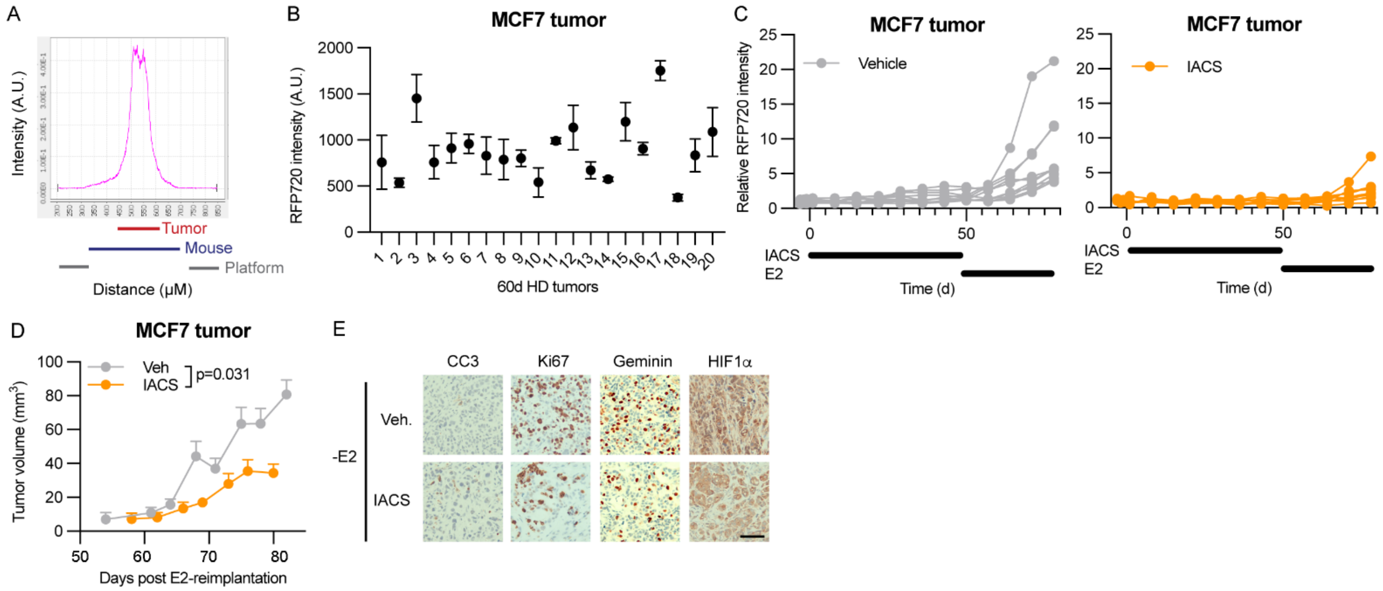

**Supplemental Figure 8: OXPHOS inhibition synergizes with endocrine therapy in PDX models of ER+ breast cancer.** (A/B) Individual tumor volume curves of HCI-017 (A) and HCI-003 (B) xenografts treated with vehicle, fulvestrant, IACS, or the combination. (C/E) Representative IHC images of HCI-017 (C) and HCI-003 (E) tumors harvested at the end of the study. Scale bar = 50 um. (D/F) Quantification of IHC in (C/E). Data were analyzed by ANOVA with Bonferroni multiple comparison-adjusted posthoc test.

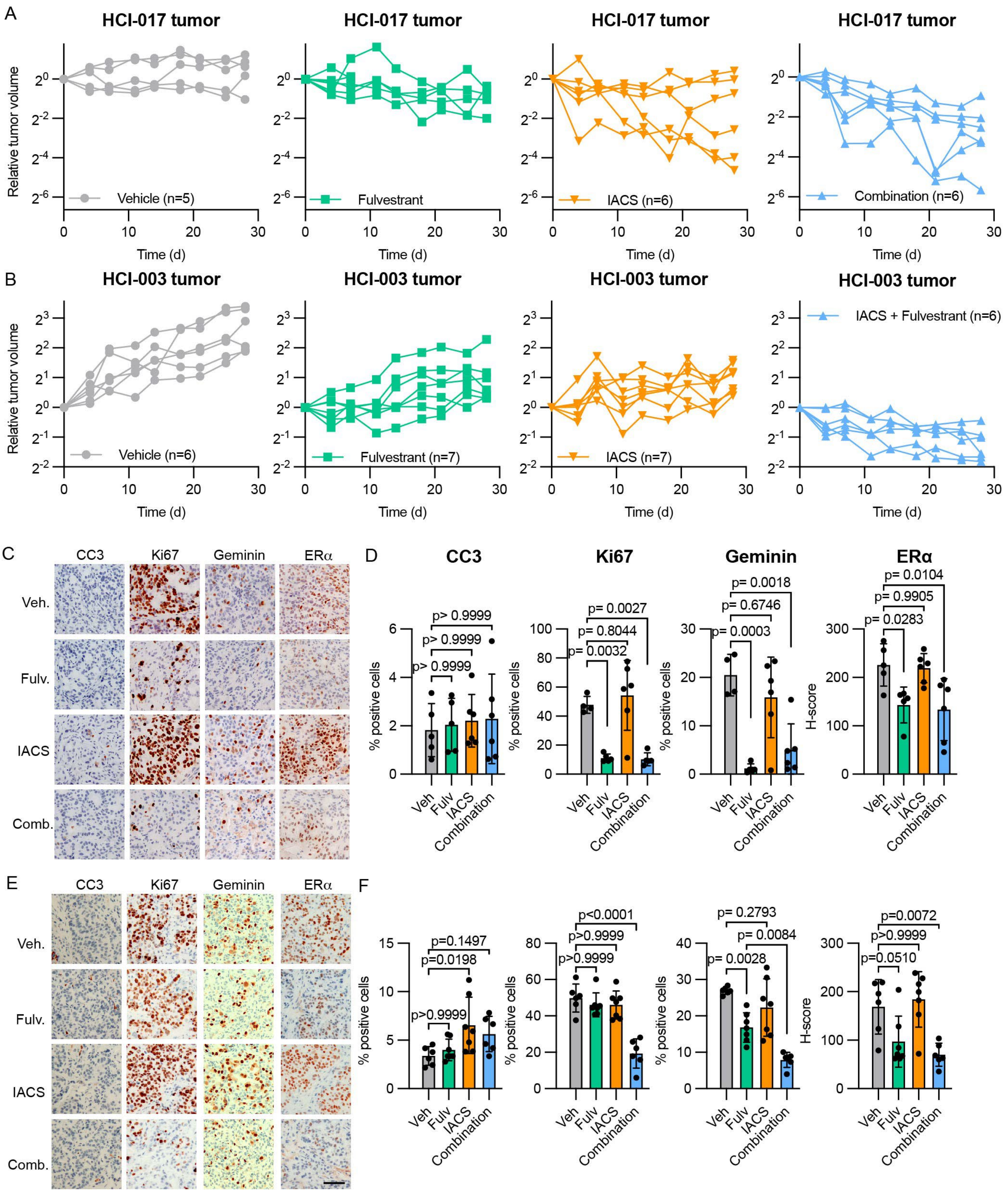
A
HCI-017 tumor
Relative tumor volume
Vehicle (n=5)
Fulvestrant
IACS (n=6)
Combination (n=6)
Time (d)
B
HCI-003 tumor
Vehicle (n=6)
Fulvestrant (n=7)
IACS (n=7)
IACS + Fulvestrant (n=6)
C
CC3
Ki67
Geminin
ERα
Veh.
Fulv.
IACS
Comb.
D
CC3
Ki67
Geminin
ERα
% positive cells
H-score
p> 0.9999
p> 0.9999
p> 0.9999
p= 0.0027
p= 0.8044
p= 0.0032
p= 0.0018
p= 0.6746
p= 0.0003
p= 0.0104
p= 0.9905
p= 0.0283
Veh
Fulv
IACS
Combination
E
CC3
Ki67
Geminin
ERα
Veh.
Fulv.
IACS
Comb.
F
p=0.1497
p=0.0198
p>0.9999
p<0.0001
p>0.9999
p>0.9999
p= 0.2793
p= 0.0084
p= 0.0028
p=0.0072
p>0.9999
p=0.0510
Veh
Fulv
IACS
Combination
